# Hippocampal sharp wave ripples and coincident cortical ripples orchestrate human semantic networks

**DOI:** 10.1101/2024.04.10.588795

**Authors:** Akash Mishra, Serdar Akkol, Elizabeth Espinal, Noah Markowitz, Gelana Tostaeva, Elisabeth Freund, Ashesh D. Mehta, Stephan Bickel

**Author notes:** Corresponding authors: Akash Mishra and Stephan Bickel.

## Abstract

Episodic memory function is predicated upon the precise coordination between the hippocampus and widespread cortical regions. However, our understanding of the neural mechanisms involved in this process is incomplete. In this study, human subjects undergoing intracranial electroencephalography (iEEG) monitoring performed a list learning task. We show sharp-wave ripple (SWR)-locked reactivation of specific semantic processing regions during free recall. This cortical activation consists of both broadband high frequency (non-oscillatory) and cortical ripple (oscillatory) activity. SWRs and cortical ripples in the anterior temporal lobe, a major semantic hub, co-occur and increase in rate prior to recall. Coincident hippocampal-ATL ripples are associated with a greater increase in cortical reactivation, show specificity in location based on recall content, and are preceded by cortical theta oscillations. These findings may represent a reactivation of hippocampus and cortical semantic regions orchestrated by an interplay between hippocampal SWRs, cortical ripples, and theta oscillations.

## Introduction

Human memory depends on an intricate bidirectional interplay between the hippocampus and memory-containing cortical regions. There is great interest in understanding the mechanisms by which these distant regions communicate because they may elucidate the processes that underlie human memory and represent potential therapeutic targets for the amelioration of memory impairment. Hippocampal sharp-wave ripples (SWR) are high frequency (80-140 Hz) oscillations that reflect coordinated neuronal population firing in the hippocampus and are associated with the hippocampal recruitment of cortical regions during memory encoding and recall^1,2^. SWRs may play a role in a two-stage framework of memory where information is encoded in the hippocampus and consolidated in the cortex, and during memory retrieval, the hippocampus reactivates these memory-containing cortical sites^3–10^. Recent studies have supported this theory by demonstrating human SWR-associated increases in high frequency activity (HFA, 70-150Hz) of specific memory-containing cortical regions during memory encoding and recall^7,8,11–13^. Intriguingly, other studies have explored ripple-like oscillatory phenomena in cortical regions (“cortical ripples”) that are temporally-coincident and morphologically-similar (frequency and duration) to SWRs^9,14–19^. However, the connection between these phenomena and memory performance remains unclear. This includes whether SWR-coincident cortical ripples show specificity for region or content and how oscillatory cortical ripples differ from non-oscillatory broadband high frequency activity (BHA), which is typically associated with neuronal population activation^20–22^. Finally, as coincident ripples reflect periods of heightened hippocampal-cortical binding^15^, there is a potential overlap in function between human ripple oscillations and theta oscillations. A potential relationship between these two has been demonstrated in human NREM sleep^23^, but this remains to be further explored in the awake state.

The semantic processing network contains precisely-coordinated cortical regions that are activated during speech perception^24–27^. This network reflects an intricate hierarchical processing ensemble that contains the primary auditory cortex and progresses to recruit widespread cortical regions. The level of comprehension and complexity increases as higher-order regions are recruited (e.g. syllables to words to sentences)^28–30^. Recent studies have identified this large, distributed cortical network to involve frontal, anterior and posterior temporal, and parietal regions^26,27,31^. Further, the anterior temporal lobe (ATL) may serve as a critical node in this network, as it coordinates semantic processing^32–35^ and serves as the interface between the posterior temporal lexical-semantic areas and frontal cortex regions^36^. This warrants the investigation of whether SWRs show specific, functional interaction with this network during word list memory encoding and retrieval.

We used an auditory word list learning task with verbal free recall to examine the relationship between ripples (SWRs and cortical ripples) and the activation of semantic networks during encoding and recall. We recorded intracranial electroencephalography (iEEG) from individuals undergoing invasive neuromonitoring for the treatment of intractable epilepsy to gain rare neural recordings of memory and semantic networks with high spatial and temporal resolution. We show that SWRs are associated with both BHA and cortical ripples in specific cortical semantic network regions. Further, coincident SWR-ATL cortical ripple oscillations are preceded by ATL cortical theta oscillations. These results extend our understanding of SWRs to the dynamic activation of semantic networks and elucidate the mechanisms by which SWRs and cortical ripples coordinate the hippocampal-cortical memory network.

## Results

### Behavioral results

Nine participants performed a word list learning task with verbal free recall (Figure 1A; Figure S1). On average, subjects recalled 4.1±2.0 words (of 12 total) per trial across 1.6±0.8 recall events (defined as recall onsets with an inter-response interval exceeding 4 seconds) per trial (median recall duration was 2.4 seconds). Participants showed a trend towards recalling more words in trials where words were assembled into sentences (sentence condition) compared to when they were not (non-sentence condition) (Z=1.953, p=0.051).

**Figure 1.**
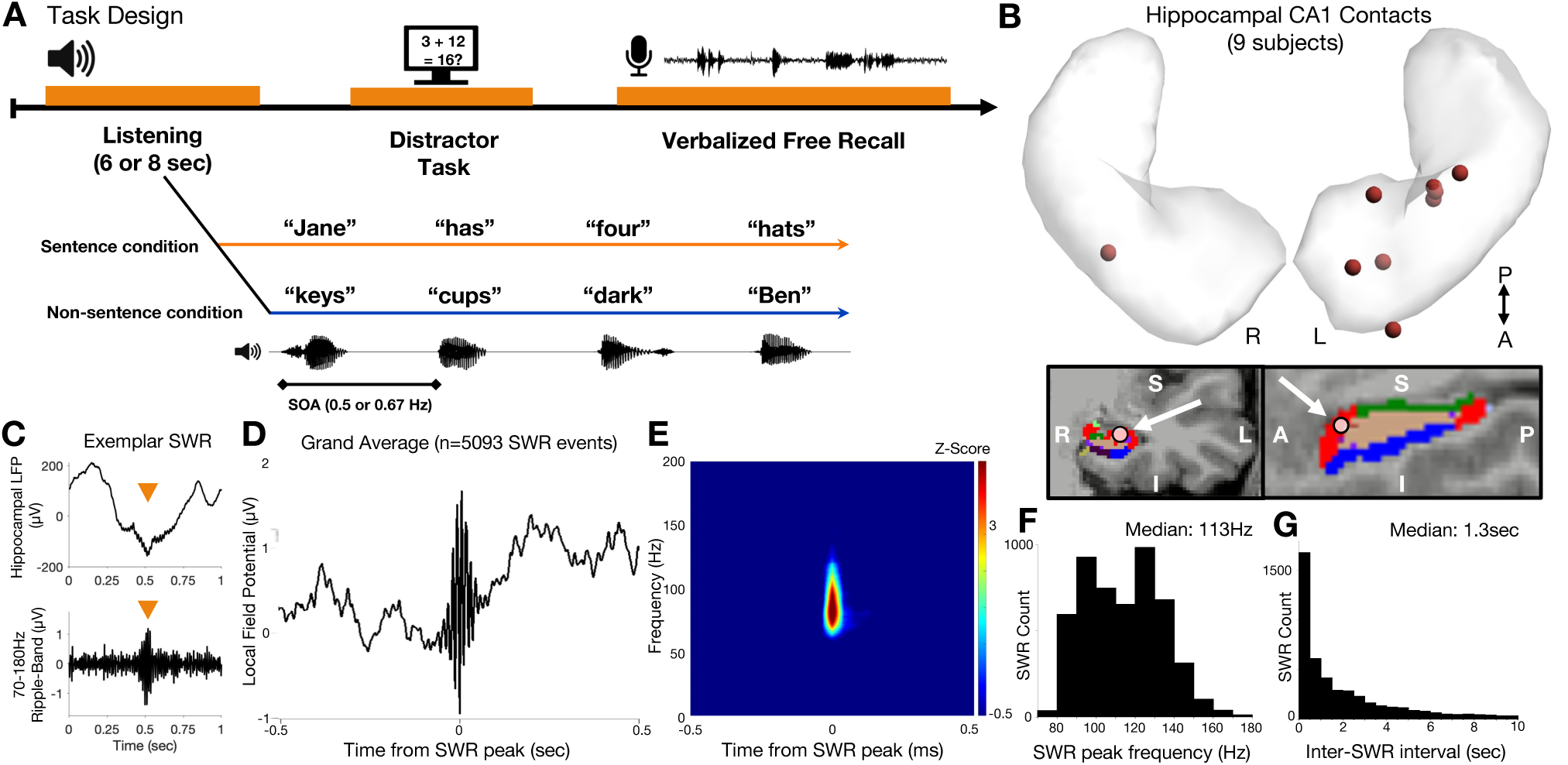
Experimental design and hippocampal sharp wave ripple detection. (A) Experimental design and example stimuli presented to subjects. Subjects listened to a list of twelve words assembled into either 3 sentences (of 4 words each) or a randomized word list. Then, after an arithmetic distractor task, subjects were asked to freely recall as many words as possible. Sentence and non-sentence trials were randomized and interleaved for a total of 30 trials. (B) (Top) Hippocampal depth electrode selection across all subjects. One point represents an individual subject. (Bottom) Representative structural reconstruction of hippocampal depth electrodes in one patient in coronal (left) and sagittal (right) views; white arrows denote CA1 recording site implemented for SWR detection analysis. Red parcellation indicates CA1 subregion. (See Figure S2 for similar reconstructions for other subjects). (C) Example of SWRs as they appear in (*top*) the raw hippocampal local field potential and (*bottom*) the 70-180 Hz ripple-band. (D) Grand average peri-SWR field potential locked to SWR peak for n=5093 SWRs from 9 subjects. (E) Wavelet spectrogram locked to SWR peak for n=885 SWRs from one representative patient. Warmer colors indicate a higher spectral power in that frequency-time combination. (F) Distribution of SWR peak frequencies and (G) inter-SWR interval for all n=5093 SWRs.

### SWR rate is modulated by encoding of word lists and subsequent free recall

We identified SWRs from the local field potentials (LFP) of hippocampal contacts located within (or as close as possible to) the CA1 subfield (Figure 1C; Figure S2). Across 9 subjects, we identified 5093 SWRs during all task periods (encoding, distractor, and recall) (Figure 1D-1E). The median peak SWR frequency across all subjects was 113Hz (Figure 1F) and the median inter-SWR interval duration was 1.3sec (Figure 1G).

Since SWRs play a role in both encoding and recall^1^, we investigated the SWR rate during these task periods. First, we created peri-event time histograms (PETH) of SWRs during the list presentation (n=7 subjects) (Figure 2A). Two subjects were excluded from this analysis, as they performed the task with a different inter-stimulus interval duration (500ms instead of 667ms; PETH for these two subjects is shown in Figure S3). After removal of 33 trials where external interruption or misclick occurred, we assessed 177 word presentation trials (8-second word list presentation, followed by a 2-second rest period). The SWR rate increased starting 1400 ms after presentation onset and decreased 800ms after presentation offset (p<0.001, permutation test) (Figure 2A). SWR rate did not significantly differ for presentation of sentence and non-sentence word lists (Figure S4). Next, we assessed SWR dynamics during memory retrieval by generating a PETH locked to verbal free recall onsets. SWR rate was transiently increased 1000ms prior to recall onset (n=342 trials; p=0.037, permutation test) (Figure 2B). Importantly, SWR rate was not reliably modulated by vocalizations captured during the task period, similar to a previous study^8^ (Figure S5).

**Figure 2.**
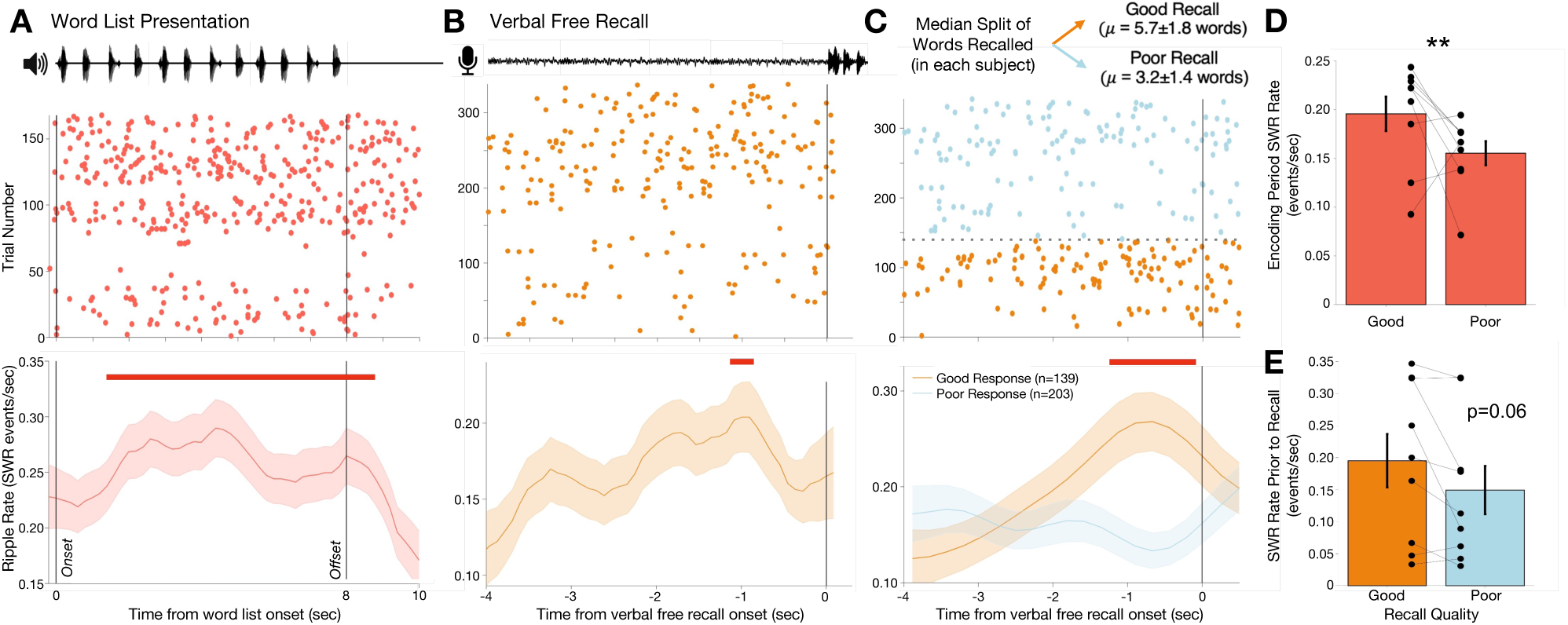
Sharp wave ripple dynamics during word list presentation and free recall. (A) SWR raster plot and peri-event time histogram (PETH) time-locked to the onset of word list presentation (n=168 trials across 7 subjects), depicting a sustained increase in SWR rate in response to word list presentation compared to baseline that subsequently diminishes after offset of word presentation (vertical line). Two subjects were removed from this analysis due to variation in word presentation frequency but demonstrated a similar SWR increase (Figure S3). Red line indicates significance at p<0.05 (permutation test compared to shuffled SWR times in the 2-second post-trial resting period). Shaded areas represent one bootstrap standard error computed over SWR events. (B) SWR raster plot and PETH time-locked to the onset of distinct recall vocalizations (n=338 trials across 9 subjects), indicating a transient response-locked increase in SWR rate. Red line indicates significance at p<0.05 (permutation test shuffling SWR times across this PETH epoch). Shaded areas represent one bootstrap standard error computed over SWR events. (C) SWR raster plot and PETH time-locked to onset of distinct recall vocalizations. Data is median split by the number of words recalled during that trial where “good response” (n=139 trials) was a trial where the patient recalled more than their median number of recalled words across trials and a “poor response” was equal to or below the median (n=203 trials). Red line denotes significance at p<0.05 (cluster-based permutation test clustering across time, p<0.05). Shaded areas represent one bootstrap standard error computed over SWR events. (D) Mean SWR rate during the encoding phase for trials that contained a good response and poor response. Each point represents the mean SWR rate per subject and lines connect data for each subject. Error bar depicts standard deviation across subjects. Stars denote significance at p<0.05 (Mann-Whitney U). (E) Mean SWR rate during the two second window prior to onset of verbal free recall in trials that contained a good response and poor response. Each point represents the mean SWR rate per subject and lines connect data for each subject. Error bar depicts standard deviation across subjects.

As SWRs represent a potential marker for effective memory processes, increased SWR rates during memory encoding and recall may be associated with improved memory performance. We separated trials into two groups based on whether subjects executed a good (number of words recalled in the trial was above the patient’s median across all trials) or poor response (below or equal to the median). Across subjects, the mean number of words recalled during a good recall was 5.7±1.8 words (139 events) compared to 3.2±1.4 for poor recalls (203 events). We then compared SWR dynamics prior to recall onset for good and poor responses. SWR rate was higher for recalls that corresponded to good response trials in the 1000ms window prior to verbal free recall compared to poor responses (p=0.0314, permutation test) (Figure 2C). Finally, we examined the relationship between SWR rate during encoding/recall on memory performance on the group level (one rate value per subject). SWR rate during the word list presentation period was higher for good response trials (0.222 SWRs/second) compared to poor response (0.167 SWRs/second) (two-tailed Mann-Whitney U=109, n1=n2=9, p=0.0379) (Figure 2D). This group-level relationship holds a trend when assessing SWR rate in the window around free recall onset (−2 to +0.5 seconds relative to free recall onset; two-tailed Mann-Whitney U=39, n1=n2=9, p=0.0547) (Figure 2E). Therefore, SWRs may reflect effective memory network activity during word list encoding and retrieval.

### SWRs reactivate functional semantic network regions during free recall

SWRs are thought to facilitate the interaction between the hippocampus and specific cortical regions^1^. Hence, SWRs during the recall portion of a word list learning task may reactivate cortical regions that were engaged during word list encoding. We first defined cortical semantic networks by assessing HFA during word list presentation in all cortical contacts (Figure S1), similar to a recent human iEEG study^31^. We identified 150 contacts that exhibited higher HFA during the listening of sentence compared to non-sentence word lists (“sentence-responsive”) and 79 cortical contacts that showed the opposite HFA profile (“non-sentence-responsive”) (FDR-corrected p<0.01, permutation test). Figure S6 depicts the difference in HFA activation between preferred and non-preferred trials across all contacts. Identified contacts were distributed across several higher-order cortical regions, further supporting the presence of a large cortical semantic network (Figure 3A).

**Figure 3.**
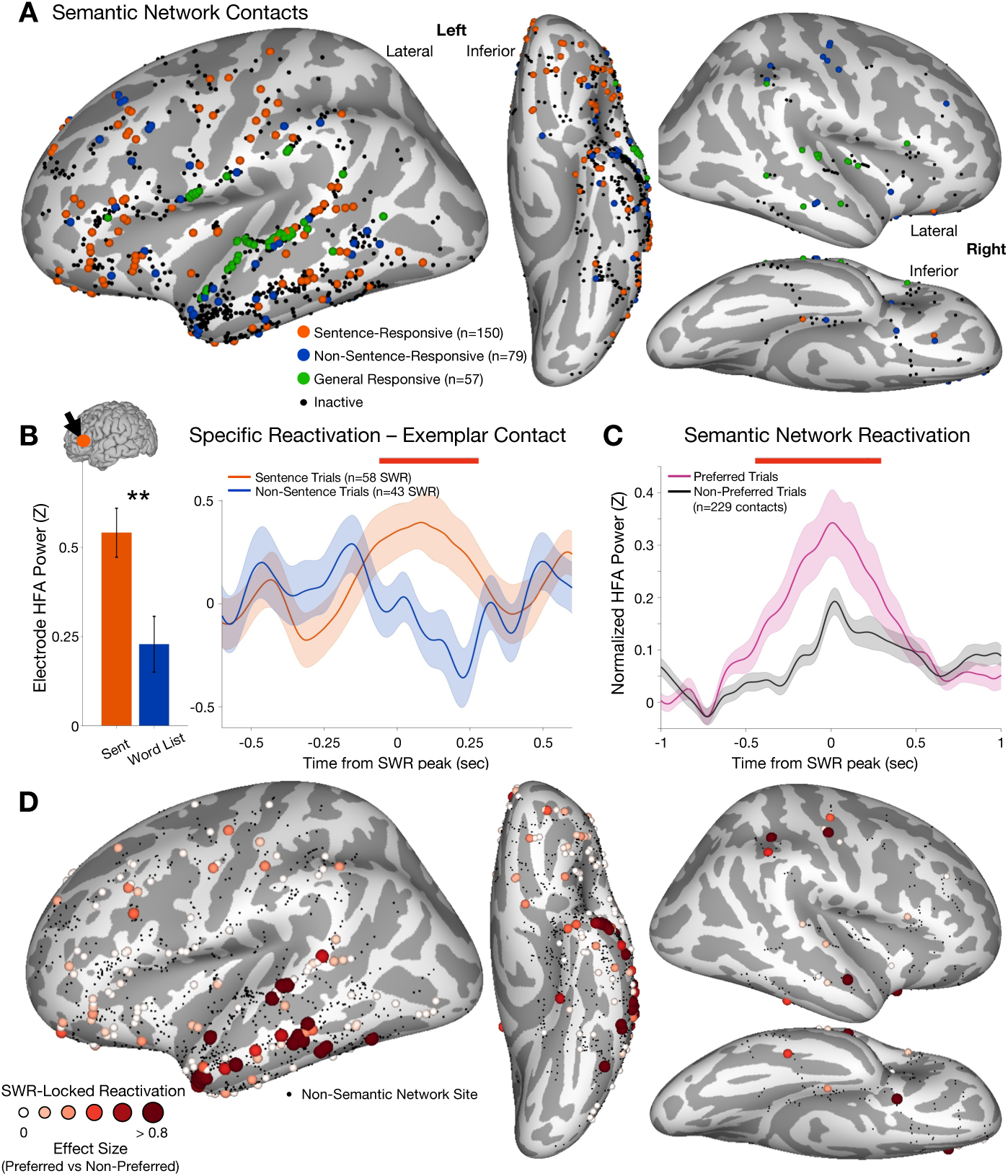
Sharp wave ripple-locked reactivation of the semantic network. (A) Left lateral *(left)*, left inferior *(middle)*, right lateral *(right upper)*, and right inferior *(right lower)* views on inflated brains of locations of sentence-responsive (blue, n=150), non-sentence-responsive (blue, n=79), and general semantic-responsive (green, n=57) contacts on an inflated brain. (Also see increase in HFA during preferential word list presentation in Figure S6.) (B) Peri-SWR HFA responses in an exemplar electrode. *(Left)* HFA response during word presentation by sentence and non-sentence trial (n=15 trials and n=180 words each, error bars denote standard deviation across trials, stars denote significance at p<0.05, two-sample t-test), *(inset)* location of the selected electrode, *(right)* HFA response locked to SWRs that occurred in the recall period when subjects recalled words from either sentence (n=58 SWRs) or non-sentence trials (n=43 SWRs). Red bar represents significant time bins at p<0.05 (cluster-based permutation test clustering across time). Shaded areas represent one standard error computed over SWR events. (C) HFA time-locked to peak of SWR events that occurred during recall of trials that aligned with the contact’s preference as compared to when they did not. Red line denotes significant time bins at p<0.05 (cluster based permutation test, n=212 bipoles). Shaded areas represent one standard error computed across contacts. (Also see spectrogram of this analysis in Figure S8.) (D) Reactivation effect size (in Cohen’s d) for semantic network contacts (combining sentence-responsive and non-sentence-responsive contacts) comparing peri-SWR HFA reactivation during the recall period of preferred versus non-preferred trials. Darker colors and larger electrode sizes depict larger effect sizes.

We then assessed the interaction between SWRs and HFA in this semantic network during recall periods. We observed an SWR-associated HFA increase in cortical contacts when there was congruence between the electrode preference and trial type (“preferred” trial-electrode combination) compared to when there was no congruence (“non-preferred” combination) (p=0.031, cluster-based permutation test) (Figure 3C). This effect is also readily visible at the single contact level (sentence-responsive contact: n=180 words in sentence list, n=168 words in non-sentence list; two-sample t(346)=3.016, p=0.027; reactivation: cluster-based permutation test clustering timebins, p=0.018) (Figure 3B). To further evaluate the spatial distribution of this effect, we calculated the effect size of SWR-locked HFA reactivation between preferred and nonpreferred trials in a 500ms window around SWR peak (Figure 3D; Figure S7; spectrograms in Figure S8). Temporal lobe contacts displayed the highest difference between preferred and non-preferred SWR-locked reactivation.

### SWR-Associated ATL HFA: Coincident Cortical Ripple or BHA?

Prior studies have shown SWR-coincident ATL cortical ripple in a word association task^9,13,37^. We similarly noted a prominent SWR-associated HFA activation in the ATL (Figure 3D) that was suspicious for cortical ripples. Hence, we aimed to replicate these findings and further investigate the relationship between non-oscillatory (BHA) and oscillatory (cortical ripple) activity in this region.

We first assessed HFA increases locked to SWRs in the recall period (500ms window centered on SWR peak) across all cortical contacts (Figure 4A; effect size in Figure S9) and identified a prominent increase in the area corresponding to the ATL. We then isolated ATL contacts (n=218) using anatomical reference and excluded 22 contacts that overlapped with the functionally-defined semantic network (Figure 4B). Not surprisingly, SWRs were associated with a significant increase in ATL HFA (p<0.001, cluster-based permutation test) (Figure 4C), maximally at the time of the SWR peak. Spectral analysis of the SWR-locked cortical activity showed a spectral peak in the 80-120 Hz range as well as an increased BHA (Figure 4D). Since the spectral peak was found at the expected frequency range of cortical ripples, we conducted a ripple detection analysis for all ATL contacts. ATL cortical ripples were found to have peak frequencies predominantly within the 80-140 Hz range (Figure 4E) but centered around 114 Hz (Figure 4F) and with a median inter-ATL cortical ripple interval of 1.9 seconds (Figure 4G).

**Figure 4.**
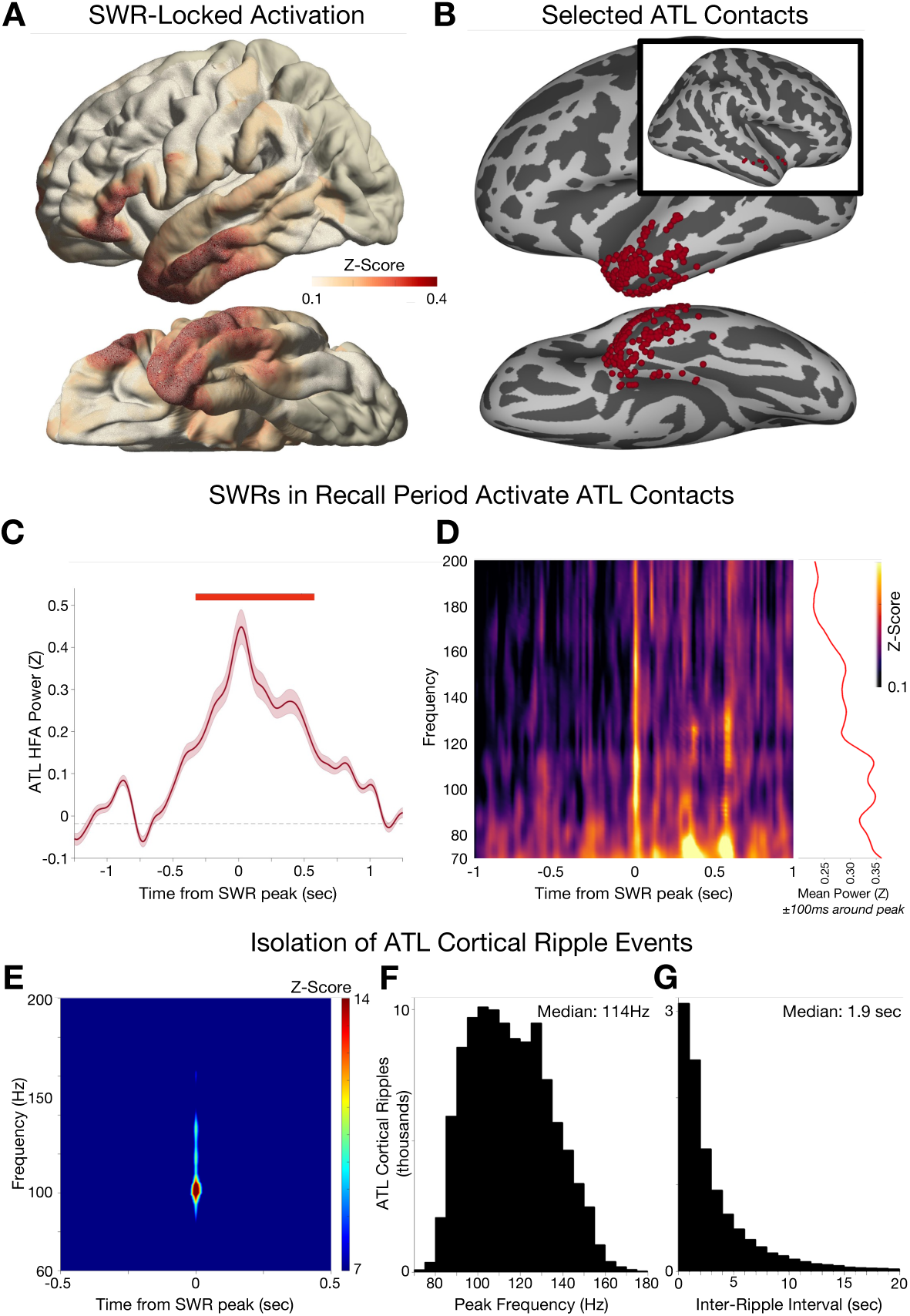
Properties of anterior temporal lobe cortical ripples. (A) Left lateral *(top)* and left inferior *(bottom)* views of normalized brains with overlaid heatmap representing regions of increased HFA increase locked to peaks of SWRs within the recall period across all contacts. Darker colors represent increased HFA, and uncolored regions represent no electrode coverage in the region. (B) Left lateral *(top)*, left inferior *(bottom)*, and right lateral *(inset)* views on inflated brains depicting the location of n=196 anterior temporal lobe contacts across subjects. Contacts were selected on patient-specific imaging and precise location may be distorted in the transfer to standardized space. (C) Increase in HFA, locked to peaks of SWRs within the recall period for n=196 ATL contacts. Red line denotes significant timebins at p<0.05 (cluster-based permutation test against pre-SWR baseline). Shaded areas represent one standard error computed across ATL contacts. (D) *(Left)* Wavelet spectrogram of ATL high frequency power locked to SWRs alongside *(Right)* mean power in the 40ms window centered around SWR peak for each frequency represented in the spectrogram depicting an increase in the 90-120 Hz range. Hotter colors represent increased power. (E) Wavelet spectrogram time locked to ATL cortical ripple peak (n=7773 events from one representative patient with n=12 ATL contacts). Warmer colors indicate a higher spectral power in that frequency-time combination. (F) Distribution of peak frequencies and (G) and inter-ripple duration for all detected ATL cortical ripples (n=108510 events in n=196 contacts across n=9 subjects).

We next assessed the coincidence of ripple oscillations by computing a cross-correlogram between SWRs during the recall period and ATL cortical ripples. There was significant coincidence (p<0.01, permutation test), with a mean delay of cortical ripple peak relative to SWR peak of 14ms (Figure 5A). Further, the proportion of SWRs that were associated with an ATL cortical ripple increased 700ms prior to verbal free recall and was sustained until recall onset (p=0.0354, permutation test) (Figure 5B). Hence, SWRs increase in the one-second window prior to recall (Figure 2B) and these SWRs are more likely to co-occur with ATL cortical ripples. Interestingly, compared to non-coincident ATL cortical ripples, coincident ATL cortical ripples in the recall period were of higher frequency (median 119 Hz compared to 113 Hz; Z=5.868, p<0.001) and shorter duration (80ms compared to 96ms; Z=7.665, p<0.001) (Figure S10).

**Figure 5.**
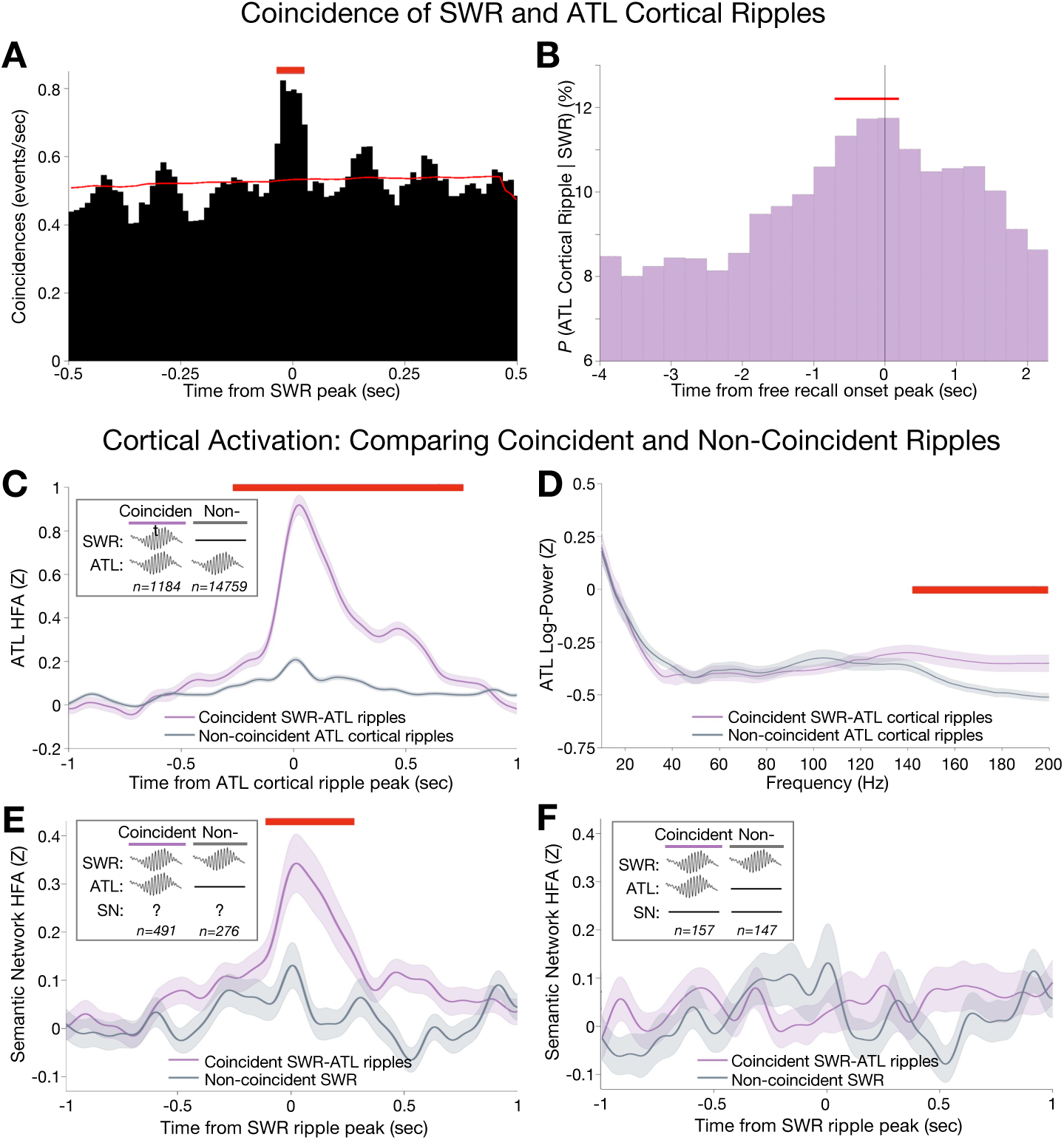
Ripples and ripple-locked activity across the hippocampus, anterior temporal lobe, and other semantic areas. (A) Cross-correlograms between ATL cortical ripples (n=15943 across n=196 ATL contacts) and SWRs (n=772 across n=9 hippocampal contacts) within the recall period across n=9 subjects. Red line denotes significant time bins at p<0.05 (permutation test with jittered cortical ripple timing), indicating high coincidence. (B) Joint probability of coincident SWR and ATL cortical ripple as a function of time relative to verbal recall onset. Red line denotes significant time bins at p<0.05 (permutation test with jittered cortical ripple timing). A significant increase is visualized within one second window prior to verbal recall onset. (C) ATL HFA locked to peak of ATL cortical ripples in the recall period separated by whether the ATL cortical ripple was coincident with an SWR (n=1184) or not (n=14759). Red line denotes significant time bins at p<0.05 (cluster-based permutation test). Shaded area represents standard error computed across ripple events. (See spectral profile of this difference in Figure S11.) (D) High frequency spectral profile of ATL cortical ripples in the recall period that were associated with SWRs compared to not coincident with SWRs. Red line denotes significant frequencies at p<0.05 (cluster-based permutation test). Shaded area represents one standard error computed across ripple events. (E) HFA in semantic network contacts, locked to peaks of SWRs in the recall period by whether the SWR was associated with an ATL cortical ripple (n=491) or not associated (n=276). Red line denotes significant time bins at p<0.05 (cluster-based permutation test). Shaded areas represent one standard error computed across ripple events. (F) Same as in (E) but with all SWR epochs containing a coincident cortical ripple within the semantic network removed. Shaded areas represent one standard error computed across ripple events.

The presence of cortical ripples raises an important methodological question: are SWR-locked increases in HFA a product of synchronous ATL cortical ripples, non-oscillatory neuronal population firing (BHA), or both? ATL cortical ripples during the recall period that were coincident with an SWR exhibited a greater increase in HFA compared to those not coincident with an SWR (p<0.001, cluster-based permutation test) (Figure 5C). Both SWR-coincident and non-coincident ATL ripples exhibited a significant increase in ripple-band power (especially between 80-140 Hz), but only coincident ATL cortical ripples exhibited a significant power increase above 140 Hz (p=0.032, cluster-based permutation test) (Figure 5D; spectrograms in Figure S11). This is suggestive of increased non-ripple related BHA associated with SWR-coincident ATL cortical ripples.

### Coincident Hippocampal-Cortical Ripples Across the Semantic Network

Coincident ripple oscillations may serve as a precise mechanism for synchronizing the activity of multiple nodes within the hippocampal-cortical memory network. Hence, we examined the impact of coincident ripples on the recruitment of cortical regions, with an emphasis on elucidating the dynamics of SWRs, cortical ripples in the ATL, and cortical ripples across the other regions of the functionally-defined semantic network (here, combining sentence-responsive and non-sentence responsive contacts). First, SWRs that were coincident with ATL ripples during the recall period exhibited a greater increase in semantic network HFA (n=491 coincident hippocampal-ATL ripples, n=276 non-coincident SWRs) (p<0.001, cluster-based permutation test) (Figure 5E). This may indicate that coincident hippocampal-ATL ripples more optimally recruit the cortical semantic network.

However, as with above, this may be a product of SWR-coincident cortical ripples in the semantic network. To investigate this possibility, we identified semantic network cortical ripples and repeated the same analysis as in Figure 5E but after removing all SWRs where there was a coincident semantic network cortical ripple. We no longer found a significant semantic network HFA increase resulting from coincident hippocampal-ATL ripples (p=0.293, cluster-based permutation test; n=157 SWRs with coincident ATL cortical ripple and n=147 SWRs without coincident ATL cortical ripple) (Figure 5F). Hence, it is probable that the observed effect of coincident hippocampal-ATL ripples on semantic network HFA is influenced by coincident semantic network cortical ripples.

We then investigated whether semantic network cortical ripples show any spatial specificity during recall. We assessed whether there was an increased coincidence of ripples for trials where the SWR and semantic network contact showed congruence in preference (i.e. a SWR that occurred during free recall of a sentence trial and coincident ripple occurring in a sentence-responsive contact, and the same for non-sentence trials/contacts) compared to when there was no congruence (i.e. SWR in a sentence trial and cortical ripples in a contact that had preference for non-sentence trials, or vice versa). The rate of triple co-occurrence of ripples in the hippocampus, ATL, and semantic network contact was increased when there was congruence in preference compared to when there was no congruence (2.1% of hippocampal-ATL-cortical contact pairs showed coincidence for congruent preference compared to 1.9% of pairs showing coincidence for non-congruent preference, X^2^(1, n=346182)=18.802, p<0.001). This is indicative of spatial specificity of cortical ripples based on the content of recall.

### Coincident SWR-ATL cortical ripples are associated with cortical theta oscillations

This analysis has primarily probed the relationship between ripples and HFA/BHA, but a key missing component is the investigation of lower frequencies. Qualitatively, a possible theta modulation of the hippocampal-ATL ripple coincidence is already notable in Figure 5A. Indeed, coincident SWR-ATL ripples (n=1184) were associated with an increase in ATL low theta power (3-5 Hz) compared to non-coincident ATL ripples (n=14759) (p=0.016, permutation test) (Figure 6A). The theta power increase is roughly coincident with the ATL cortical ripple peak (p=0.003, permutation test) (inset, Figure 6A). We then confirmed the presence of theta oscillations in single trials around the coincident ATL ripples by using the FOOOF toolbox^38^. Detected theta oscillations showed a peak frequency between 3-5 Hz (Figure 6B). Finally, to determine the temporal relationship more precisely between coincident hippocampal-cortical ripple oscillations and theta oscillations, we constructed a cross-correlogram between detected theta oscillation peaks and the peaks of ATL cortical ripples that were and were not coincident with a SWR. SWR-coincident ATL cortical ripples co-occurred with ATL theta oscillations at a higher rate compared to non-coincident ATL cortical ripples (p=0.002, cluster-based permutation test) (Figure 6C). On average, the theta oscillation peak was delayed by 109ms relative to the SWR-coincident ATL cortical ripple peak. This is suggestive of a potential relationship between coincident hippocampal-ATL ripples and ATL theta oscillations.

**Figure 6.**
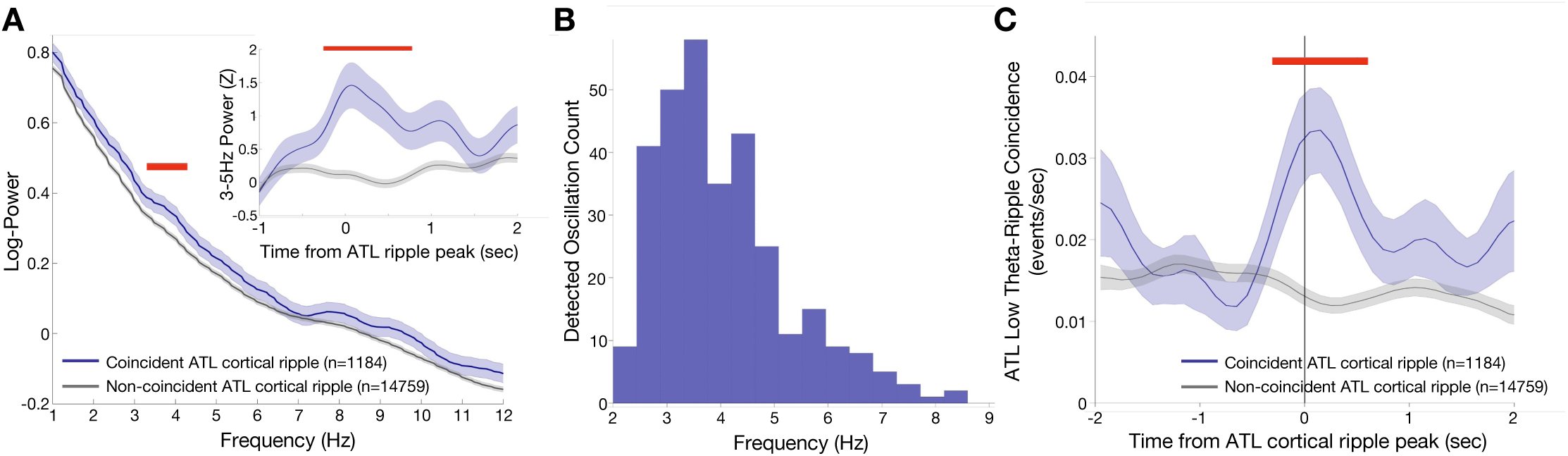
Relationship between cortical ripples and theta oscillations. (A) Low frequency spectral profile of ATL activity locked to peak of ATL cortical ripple that were coincident (n=1184) and not coincident with a SWR (n=14759). Red line denotes significant frequencies at p<0.05 (cluster-based permutation test). Shaded areas represent one standard error across ripple events. *(Inset)* Power in the 3-5 Hz low theta frequency range across time when locked to peak of ATL cortical ripples occurring in the recall period. Red line denotes significant time bins at p<0.05 (cluster-based permutation test). Shaded areas represent one standard error across ripple events. (B) Distribution of peak frequency of detected low frequency oscillations around ATL cortical ripples that were coincident with a SWR. (C) Cross-correlograms between peak of ATL cortical ripples that were coincident with SWRs during the recall period and peak of ATL theta oscillations. Red line denotes significant time bins at p<0.05 (cluster-based permutation test). Theta oscillations occurred at a peak lag of 109 ms relative to coincident ATL cortical ripples. Shaded areas represent one standard error computed across cortical theta peak events.

## Discussion

Memory formation and retrieval involves precise hippocampal-cortical coordination. Neural oscillations are a mechanism that may support this complex process. We investigated this possibility by using intracranial recordings from the human hippocampus and widespread cortical regions during a word list learning task. We show that, during encoding and recall, SWR rate increases and correlates with recall performance. During memory retrieval, SWRs exhibit reactivation of specific cortical regions based on recall content. In the ATL, a major semantic hub, SWRs are associated with both oscillatory (coincident cortical ripples) and non-oscillatory (BHA) cortical activity. Finally, coincident hippocampal-ATL ripple oscillations are more likely to be preceded by cortical theta oscillations. Taken together, we replicate prior findings in supporting a function of SWRs in the encoding and retrieval of word lists^11,39^. This supports congruence in the function and mechanisms of SWRs across memory tasks in awake humans^7–9,15,40^. Finally, we describe a complex interplay between SWRs, cortical ripples, BHA, and cortical theta in the semantic network that may underlie word list memory.

SWRs reflect highly-synchronized firing patterns that may bind activity across specific brain regions, including the hippocampus and cortex^1,41–44^. Hence, during encoding and recall, SWRs may engage multiple memory-containing cortical regions^15^. When the rate of SWRs increases during encoding, there may be increased binding of hippocampal-cortical activity that strengthens the memory representation. Similarly, during retrieval, increased SWR rate may indicate heightened reactivation of memory traces. This would manifest prior to the onset of free recall, as retrieval precedes recall. Increases in SWR rate during successful encoding and retrieval have been identified previously in awake humans^7–9,11,45^. Our findings align with these studies, as we identify increases in SWR rate during both word list presentation and prior to free recall onset. Further, for trials where more words were recalled, SWR rate was higher during both encoding and prior to recall onset.

HFA is one method by which SWR-associated cortical activity may be assessed, as it is the part of the iEEG signal that best correlates with neuronal population spiking^46,47^. HFA has been utilized previously to identify SWR-associated activity in visual processing regions^8^, the default node network^7^, and in sleep states, more dispersed cortical networks^10,12^. Here, we examined if this association extends to the semantic network. The auditory presentation of a set of words engages an assembly of interconnected cortical nodes from lower-order primary auditory cortical regions to higher-order regions involved in extracting rich semantic information^27,31^. To determine contacts located in the semantic network, we identified sentence-preferring and non-sentence-preferring contacts based on their activity during encoding. These contacts were found to be present in widespread frontal and temporal cortical regions, similar to prior studies that used a similar approach^25,27,31^. During recall, we found increased cortical HFA around SWRs when the content of the recall aligns with the functional preference of the contact. This finding extends our knowledge of SWR-associated cortical activation during recall into the semantic domain. It is interesting to note that the HFA activation begins 400ms prior to the SWR peak. This observation may support a hypothesis that the cortex initiates memory retrieval by biasing hippocampal activity to generate SWRs, which then function to reciprocally activate specific cortical memory representations^4–6,43,44,48,49^. However, to confirm this, further studies are needed, ideally by including single-or multi-unit recordings.

We next expanded our analysis beyond this functionally-defined semantic network to the ATL. This was motivated by recent human word association studies suggesting a role of SWRs in the reactivation of specific neuronal assemblies in this region^37^. Furthermore, the ATL was one of the most activated areas in our data-driven SWR-locked cortical HFA analysis (Figure 4A; Figure S9). While the exact function of the ATL is still largely unknown, the hub-and-spoke theory suggests that this region is an orchestrator of higher-order semantic network cortical regions. The ATL is thought to facilitate the activation of individual semantic processing nodes to effectively retrieve higher-order semantic concepts^50,51^. This aligns with clinical outcomes where patients with anterior temporal lobectomies of the dominant cortex exhibit increased risk of naming deficits^52–54^. We found that SWRs during the recall period are associated with coincident increases in ATL HFA. This increase consists of both BHA and a spectral peak in the range of previously-reported cortical ripples. Cortical ripples are often temporally-coincident with SWRs and may play a key function in rodent^19^ and human memory^9,13,15,37,55^. These events may signify the precise binding of SWRs to the reinstatement of cortical memory representations^15,16^. In support of this hypothesis in the domain of verbal memory, studies have utilized a word association task to demonstrate the coincidence of temporal cortex ripples with SWRs, and have shown cortical ripples to contain specific neuronal firing patterns indicative of memory representation activation^9,13,37^. More recent studies have found this phenomenon to be more ubiquitous across the human cortex, with specific hippocampal-cortical ripple assemblies reactivated during memory retrieval^15,16^. These findings partially parallel rodent studies that demonstrate SWR-coincident cortical ripples in association cortices^19^.

We identified cortical ripples in the ATL, and 7.4% of them exhibited coincidence with a SWR during the recall period. This is similar to a rodent study that reported 20% of association cortex ripples co-occurring with SWRs^19^. There may be several factors that drive this scarcity: (1) poor proximity of recording contacts to ripple-generating nodes^56^; (2) suboptimal detection parameters for SWRs and cortical ripples^57^; (3) recall events may not require coordination of hippocampal-cortical networks^58^; and/or (4) co-ripple rates are lower in humans compared to rodents. Furthermore, cortical ripples were of similar duration and frequency compared to SWRs. However, SWR-coincident ATL cortical ripples were of higher frequency and shorter duration compared to when not SWR-coincident. We speculate that this may be due to different subtypes of cortical ripples. Long-duration ripples are associated with improved memory consolidation^6,59,60^, but shorter-duration ripples may contain more succinct neuronal firing that communicate smaller packets of information. It is also possible that inter-areal communication via coincident ripples requires similarity in frequency and/or duration. Next, we find that, even when the increase in SWRs prior to recall is taken into account, the proportion of coincident ripples increases prior to recall. Hence, ATL cortical ripples are not exclusive to periods of SWRs but their coincidence increases in periods of memory retrieval. To add to this, coincident hippocampal-cortical ripples are more likely to occur in regions that were previously engaged during content encoding, indicating that the content of recall influences the presence of cortical ripples in the semantic network. These findings suggest that cortical ripple oscillation networks exist independently of SWRs, but coincidence of SWRs and cortical ripples occur in times where more optimal synchrony is required^15^. To summarize, coincident hippocampal-cortical ripple oscillations display unique characteristics that may support heightened memory network activity. One critical piece of this puzzle that is still missing is the directionality or causality of this relationship. Intriguingly, it is possible that the hippocampus itself is not the primary (or “driving”) oscillator. This “cortical-centric” approach strays from prior human ripple oscillation studies, which have predominantly focused on SWRs. This may also point towards a more general function of cortical ripples in human cognition outside of memory.

The human SWR literature contains reference to both non-oscillatory (BHA)^7,8^ and oscillatory (ripple) cortical activity^9,15,16^ associated with SWRs during memory recall periods. We investigated the possibility of a co-occurrence between these two phenomena. We show that SWR-coincident ATL cortical ripples exhibit an increase in BHA, indicative of heightened non-oscillatory neuronal activity in the ATL. This feature is less prominent for ATL cortical ripples not coincident with SWRs. Hence, SWRs are associated with both oscillatory and non-oscillatory ATL cortical activity. It is possible that these mechanisms function in concert to reflect periods of heightened hippocampal-cortical network activation. Further, it is interesting that cortical ripples in the semantic network do not display a similar increase in non-oscillatory activity as in the ATL. We posit that this may be due to the relationship between cortical ripples and “backbone” spiking sequences^13,37,55^, where small variation in similar sequences alters content retrieval. Cortical ripples may re-engage the same assemblies but display slight deviations in spiking sequence. This is not easily clarified in our recording modality, but may partially explain why semantic network cortical ripples do not display increases in BHA when they are coupled with SWRs. Further, Khodagholy et al. reported that cortical ripple oscillations were not prevalent in the rodent cortex outside of association areas^19^; hence, it is possible that cortical ripples that are detected in association areas are functionally different from those in other areas. It is possible that this relates to the increase in connectivity (hippocampal-cortical and cortico-cortical) that is present in association cortices^61^.

Beyond ripple oscillations, theta oscillations have been described widely in the literature as another mechanism underlying hippocampal-cortical coordination during memory processes^62^. While SWRs mainly occur in non-theta states, evidence from rodent models suggests a relationship between other high-frequency oscillations in the epsilon band^63,64^ and theta in the hippocampus^62,65–67^ and cortex^68^.

Furthermore, a recent study investigating SWRs in humans during NREM sleep described that cortical theta oscillations temporally precede SWRs in some regions, while they succeed them in others^23^. In our study, coincident hippocampal-ATL ripples were more likely to be associated with cortical theta oscillations in the ATL compared to non-coincident ripples, and the theta oscillation peak succeeded the coincident ripple by 109ms. This aligns with a recent study where several human temporal cortex contacts exhibited a theta-ripple relationship in NREM sleep^23^. This represents an interesting ruffle in the exploration of the mechanisms that underlie memory, as it is believed that theta oscillations underlie active memory function^62^ while ripple oscillations represent offline memory function^1^. In the awake human, these two mechanisms may function in concert such that cortical theta-ripple assemblies (even if temporally-asynchronous) may coordinate the precise reactivation of the ATL during word list retrieval. Alternatively, it is possible that concerted theta oscillations are underlying core encoding processes and coincident ripple oscillations function as an immediate replay mechanism for heightened memory consolidation. In this case, it is possible that SWRs which “successfully” co-occur with a cortical ripple may be directed by a preceding cortical theta oscillation, which more optimally engages the memory network as compared to non-coincident ripples. Finally, it is possible that current approaches of detecting human SWRs^57^ may, at least in part, detect epsilon oscillations, which would be expected to be modulated by theta oscillations^63,66,69–71^. However, that we only found co-ripples to be associated with cortical theta, and not hippocampal theta, would speak against this.

In conclusion, ripple oscillations serve as a dynamic mechanism underlying human memory function. In a word list learning task, we identify SWR-associated reactivation of specific cortical regions during free recall. There exists a unique interplay between the hippocampus and the ATL where, in times of SWRs, the ATL exhibits synchronous cortical ripple and BHA increases. Coincident ripples across the hippocampus, ATL, and semantic processing areas show specificity based on recall content and may serve as a powerful mechanism for recruitment of semantic memory-containing regions. Further, coincident ripples between the hippocampus and the ATL are more likely to be associated with ATL cortical theta oscillations. These findings extend our understanding of word memory retrieval, further elucidate the function of the ATL in coordinating activation of semantic processing regions, and provide evidence for coincident ripples as a mechanism of multi-nodal synchrony in human memory networks.

## Methods

### Subjects

Intracranial recordings were obtained from nine patients (7 females, median age 32, age range 19-58) with medically intractable epilepsy undergoing iEEG recording at Northwell Health (New York, USA) to assist in the identification of epileptogenic zones for potential surgical treatment. Patients undergo continuous iEEG monitoring for a period of 1-3 weeks, during which they may participate in cognitive and functional testing. The decision to implant, location of implanted electrodes, and the duration of implantation were made entirely on clinical grounds by the treatment team. Subjects were selected to participate in this study based on presence of adequate hippocampal and cortical coverage and ability to perform a word list learning task. The study was conducted in accordance with the Institutional Review Board at The Feinstein Institutes for Medical Research (Northwell Health), and informed consent was obtained prior to testing.

No clinical seizures occurred during or within the two-hour period prior to the experimental blocks. All participants performed the task in their primary language (English or Spanish) and all participants were found to have left-hemisphere dominance for language identified via Wada testing and language fMRI. Patient information, including number of contacts, region of seizure focus, primary language, and post-implantation treatment are provided in Table 1.

### Stimuli and Task

Construction of word sets: In each trial, twelve monosyllabic English words were selected from a set of words such that they formed a set of three sentences of four syllables each (sentence trials) or a randomized list (non-sentence trials). Sentences had the same syntactic structure: noun + verb + adjective/descriptor + noun, but each individual word did not have any relation to other presented words (i.e. there was no syntactic dependence among words and subjects could not reasonably predict the upcoming word)^72^. Fifteen sentence trials and fifteen non-sentence trials were randomly shuffled to create the thirty trials that composed the experiment. In each trial, the words were presented in an isochronous manner without any acoustic gap between them. Word stimulus onset asynchrony (SOA) was 667ms (1.5 Hz word presentation) for all subjects except subjects 1 and 2 where the SOA was 500ms (2 Hz word presentation).

Subjects were provided with instructions for the task, including a sample set of words, and we adjusted speaker volume for comfort as needed. Each trial started with a subject-initiated button press, and participants waited 500ms before being presented with either a sentence trial or a non-sentence trial. Upon the presentation of the words, participants waited 2000ms before being presented with a distractor task. In the arithmetic distractor task, participants were asked to determine whether a given arithmetic operation was true or false (e.g. “2+94=96” is true and “3+39=41” is false). The operation was visually presented one number or symbol at a time, each presented for 500ms with an inter-trial interval of 400ms. After the final number, subjects responded with a button press. In each trial, participants were presented two equations in succession. Following this, participants viewed a cue screen where they were asked to freely recall as many words as they could from the twelve words that were presented earlier in the trial. Participants pushed a button to designate that they were done recalling and were ready for the next set of words (in the next trial) to begin. For subject 3, immediately after the presentation of words, we asked the participant to recall as many words as they could in order to establish adequate comprehension of words. If the participant was unable to verbally recall any of the words from the presented set, we manually aborted the trial and the next trial began. If the participant was able to recall any words, the trial continued. Two out of thirty trials were manually aborted.

We executed this task using the Psychophysics Toolbox in MATLAB (MathWorks Inc., Natick, MA, USA). Visuals were presented using a 15” laptop or 24” monitor with sound stimuli sampled at 44.1k Hz. Sound stimuli were presented with a speaker located directly in front of the participant. To minimize distractions and external noise, all other electronic devices in the study room were switched off, we kept the patient room door closed, and clinical staff were instructed to try to not enter or exit the room during the duration of the task.

### Identification of verbal recall events

A microphone affixed to a nearby stable surface recorded the entire experimental block, including the free recall periods for each trial. The onsets and offsets of each recall event and the contents of the recall were extracted in an offline analysis using Audacity auditory presentation software (Audacity, Oak Park, MI, USA). Recall events were designated as any relevant words that the patient stated, and we marked any recall onset that occurred longer than 4 seconds after the offset of the previous recall event as an independent event. Any trials induced by a mis-click or that included outside interference were removed from subsequent analysis.

To quantify the number of recalled words, we performed a median split by patient across all trials. Based on whether the patient was able to recall more than or less than the median, we marked each trial as “effective” recall or “poor” recall, respectively. This aimed to normalize for variation in the number of recalled words per patient.

### Intracranial Recording Acquisition and Preprocessing

Intracranial recording sites were subdural grids, strips, stereo-EEG depth electrodes, or a combination thereof (Ad-Tech Medical Instrument Corp., Oak Creek, WI, USA; Integra LifeSciences, Princeton, NJ, USA; PMT Corp., Chanhassen, MN). Recording sites in the subdural grids and strips were 1- or 3-mm platinum disks with 4- or 10-mm intercontact (disk center to disk center) spacing. Recording sites in depth electrodes were 2-mm platinum cylinders with 0.8mm diameter and 4.4-mm intercontact spacing. During the recordings, we re-referenced the intracranial EEG signal to a subdermal electrode or subdural strip. Neural signals were acquired using a Tucker-Davis Technologies (TDT) PZ5M module (Tucker-Davis Technologies Inc., Alachua, FL) at either 1.5- or 3k Hz and saved for offline analysis. Prior to experimentation, signal quality and power spectra were inspected online using TDT Synapse oscilloscope software, and if needed, changes were made to improve signal quality. During the task, transistor-transistor logic pulses triggered by the stimulus presentation software were generated at specific experimental timepoints of interest (onset of each trial, onset of the distractor task, and onset of free recall cue). This served to synchronize intracranial recordings to the task-related events of interest.

We performed all data analysis in MATLAB using FieldTrip^73^ and custom written scripts. We resampled neural data to 500 Hz and we removed potential 60 Hz (and its 120 and 180 Hz harmonics) power-line noise using a notch filter (zero-lag linear-phase Hamming-window FIR band-stop filter). We visually inspected raw iEEG data to detect noisy or bad channels, which were excluded from further analysis. Then, we average referenced neural data to remove global artifacts. Seizure onset channels identified by the clinical team were excluded from analysis.

### Electrode Registration and Localization

For each participant, a pre-implant T1w structural MRI and post-implant CT scan were collected. We performed intracranial electrode localization using the iELVis toolbox^74^, which utilizes BioImage Suite^75^, FreeSurfer^76^, and custom written code. To summarize the electrode localization procedure, electrode contacts were manually registered in the post-implantation CT scan using BioImage Suite, which is co-registered to the pre-implantation MRI scan to minimize localization error due to brain shift. We used FreeSurfer to align the patient’s pre-implantation MRI to a standard coordinate space, segment the cortical surface, assign anatomical locations for each contact, and determine hippocampal subfields in each patient. Next, we used iELVis software to project the contact locations onto FreeSurfer’s standard surface. We visually inspected the location of these contacts on the subject’s brain to confirm contact location.

### Selection of Semantic Network Contacts

We determined electrode preference based on whether HFA increased significantly during the presentation window. We followed the methodology implemented by Fedorenko et al. in a previous study^31^. Sentence-responsive contacts were defined as contacts in which the magnitude of HFA was significantly higher for trials in the sentence condition compared to the non-sentence condition; contacts for which the effect was in the opposite direction were classified as non-sentence responsive. To identify the functionality and specificity in responsiveness across contacts, we constructed a 500 ms epoch of HFA for each presented word, with the epoch beginning at the onset of each presented word and ending 50ms prior to the onset of the next word, and we classified each epoch as being part of a sentence trial or a non-sentence trial. We then baseline corrected the signal using a 450ms pre-trial baseline. For each electrode, we then computed the mean HFA across the twelve word positions in each trial for each condition. We then performed a Spearman’s correlation between the single trial mean HFA and a vector of condition labels (sentences = 1; non-sentences = −1). To assess the significance of this correlation, we compared the resulting correlation coefficients (Spearman’s rho) against a null distribution obtained by randomly reordering the condition labels and calculating a new Spearman’s rho for 1000 iterations. Correlations that were in the top or bottom 2.5% of the distribution were denoted as significant. Significant contacts with a positive rho value (that is, a positive correlation) were marked as sentence-responsive, and significant contacts with a negative rho value were marked as non-sentence-responsive. Hence, the semantic network was divided into contacts that were preferentially sentence-responsive and contacts that were preferentially responsive to non-sentence trials. Separately, contacts that showed increases in HFA for both sentence and non-sentence trials as compared to baseline were marked as general language-responsive.

We also identified contacts that were in the anterior temporal lobe. To systematically define these contacts, we identified the axis between the most anterior tip of the temporal lobe to the most posterior aspect of the middle temporal lobe as defined by the Desikan-Killiany atlas^77^ overlaid onto each subject’s specific brain image. All contacts that were anterior to the orthogonal line constructed at one-third of the constructed temporal lobe axis were defined as being in the ATL.

### High-frequency broadband signal and related spectral analysis

We defined high frequency activity (HFA) signal as the mean normalized power of 70-150 Hz. For analyses in which spectrograms were computed or HFA activity was compared between contact types, we computed HFA power independently for each contact by: (1) bandpass filtering the iEEG signal in 10 Hz bands (i.e. 70-80 Hz, 80-90 Hz, 90-100 Hz, and so on, but excluding windows of 118-122 Hz to exclude potential power line noise) using a fourth-order Butterworth filter; (2) applying a Hilbert transform to each individual frequency band and taking the absolute value of the resulting envelope; (3) amplitude normalization, via division of the mean of the frequency band signal; (4) averaging all normalized frequency band envelopes. This procedure results in a single time series that serves as a proxy for mean neuronal activity at a given contact. This normalization procedure has been implemented previously^8,78^ and corrects for the 1/f decay in EEG power spectra while providing temporal smoothing at the higher frequencies. Prior to any analysis implementing this derived HFA signal, we visually inspected the data for artifacts. Candidate artifacts were identified as peaks 4 standard deviations above the mean HFA signal across contacts. Artifacts were then visually inspected, and if needed, contacts were marked to be excluded from any additional analysis.

Epochs of HFA signal were created as needed for time periods of interest, including ripple-locked, response time-locked, and word presentation-locked. For each, we normalized the HFA signal with a pre-event baseline (typically the 500ms pre-trial baseline or the 500ms period immediately after response cue). This allowed for the comparison of contacts within regions of interest across subjects. To compare HFA across two signals of interest, we implemented a two-tail nonparametric cluster-based test (clustering across time bins) using the fieldtrip toolbox^73^. For single-electrode examples, due to smaller sample sizes, we used a two-tail Wilcoxon signed-rank test.

To examine the effect size of HFA effects, we used a Cohen’s d statistic. We evaluated the effect size as the following: small (d=0.2), medium (d=0.5), and large (d=0.8)^79^. We implemented this metric when determining the magnitude of effect, by electrode, for (1) preferred and non-preferred analysis (averaging over a 500ms epoch around SWR peak for each) and (2) examining peri-SWR HFA compared to baseline (comparing 500ms epoch around SWR peak with a 500ms period preceding this).

### Ripple Detection

We performed an offline ripple detection. For SWR detection, we identified an electrode contact located in or adjacent to the lateral aspect of the body of the hippocampus (CA1/CA2 subfield region). The exact anatomical location of electrode selection in each patient is depicted in Figure S1. We re-referenced the signal to the closest nearby white matter contact, 8.8mm away from the selected hippocampal contact and filtered between 70 and 180 Hz using a zero-lag, linear-phase Hanning window FIR filter with 5 Hz transition band. We then applied a Hilbert transform to attain signal amplitude. As in a prior study^80^, we clipped this signal to 3 standard deviations to minimize bias that may arise from high ripple rates. The clipped signal was then squared and smoothed via a Kaiser-window FIR low-pass filter with 40 Hz cutoff. To attain a baseline for event detection, we used the mean and standard deviation of the pre-experiment resting period signal after removal of 100ms periods surrounding any manually-detected IEDs (please see below for methodology and rationale). Events from the original signal that exceeded 3 standard deviations above baseline were selected as candidate SWR events. We defined the onset and offset of each event as the timepoints where the ripple band power decreased below 2 standard deviations. Events shorter than 42ms (computed as the duration of 3 cycles of 70 Hz) and longer than 250ms were removed. Events where the peak-to-peak difference was under 200ms of one another were merged. We then aligned the SWR peak to the maximum amplitude of the ripple-band envelope.

To control for global and transient artifacts, we performed a SWR detection on the common average signal of all contacts. Any SWR events that occurred within 50ms of a common average ripple-band peak were removed. Further, to avoid inclusion of any pathological high-frequency discharges, specifically inter-ictal discharges (IEDs), we used a manual detection process. Candidate IEDs were identified as having an amplitude greater than 4 standard deviations above mean, and a peak width of less than 100ms. Of these candidates, true IEDs were manually selected based on patient-specific IED physiology. Following this, we implemented an automatic IED detection method; the raw hippocampal bipolar LFP was filtered between 25-60 Hz (using a zero-lag, linear-phase Hamming window FIR filter) and we applied a similar methodology to the ripples analysis above (rectifying, squaring, normalizing). Detected events that exceeded 5 standard deviations were marked as IEDs. SWR events occurring within 200ms of an IED event were rejected^81^. For identification of cortical ripples, we implemented the same ripple detection algorithm. Finally, we confirmed whether detected SWRs and cortical ripple events were true oscillations by using the eBOSC toolbox^82,83^. We mandated that all identified events had at least one oscillatory cycle within the ripple frequency range. We also executed this procedure for all cortical contacts analyzed, including ATL and semantic network contacts.

We primarily assessed SWRs in two main experimental periods: (a) perception of word list, during which participants listened to the word list and (b) recall, within four-second windows around the onset of independent verbal recall events (defined as verbal recall events that were at least four seconds apart). This aimed to maximize the potential that SWR events were directly involved in the encoding and recall process. SWR events that coincided with other parts of the task (i.e. perception of the word list in the next trial or distractor task) were removed from this sample.

Human SWRs have been proposed to predominantly occur in the 80-140 Hz frequency range and most studies utilized an 80-150 Hz frequency range for SWR detection^57^. However, we aimed to minimize the risk of filtering out ripple events, and hence, we used a wider frequency range of 70-180 Hz (similar to other studies^8,12,57^). Demonstration of the peak ripple frequencies show that most ripples were indeed within the 80-140 Hz range (Figure 1F).

### Spectral analysis and detection of aperiodic oscillatory activity

We then evaluated whether the increases in SWR-associated ATL HFA were increases in broadband HFA or cortical ripples. As ripples are rhythmic phenomena, if cortical ripples were indeed driving this effect, we would expect to identify an increase in oscillations detected within the ripple-band. Per prior studies, we expect the cortical ripples to have a peak frequency within the 80-120 Hz range^9,15–17^. We performed a wavelet-based frequency analysis for the 10-200 Hz frequency range (in 1-Hz steps; width 8 Hz) for each detected ATL cortical ripple. We removed 3 Hz bands centered on 60, 120, and 180 Hz to correct for line noise. We then separated all ATL cortical ripples that occurred within 100ms of a hippocampal SWR and compared the log-frequency spectrograms for cortical ripples that were coincident with a SWR and isolated cortical ripples without coincident SWRs. We implemented a cluster-based permutation test to determine significant frequency clusters.

To more closely assess the relationship between ATL cortical ripples and low-frequency oscillatory phenomena, we performed a wavelet-based frequency analysis for the 1-12 Hz frequency range (in 0.1 Hz steps; width 6 Hz) for each detected ATL cortical ripple. To visualize the changes in lower frequencies, we constructed plots depicting log-power as a function of frequency (averaging over a 500ms window around cortical ripple peak) and power as a function of time. We separated these plots between coincident and non-coincident ATL cortical ripples to assess for differences between the two. We confirmed these findings by implementing the FOOOF algorithm (version 1.0.0)^38^, which assists in the decomposition of a neural signal into a periodic and aperiodic component on the single-trial level. We detected low-frequency oscillations in the 2-10 Hz range on peri-SWR data (400ms epoch). We selected the minimum peak height to be 0.3 and the peak width limits were set to be 1-12 Hz. The resulting identified peaks were then plotted onto a histogram to depict most prominent SWR-associated low frequency oscillations.

### Detection of theta oscillations

We performed theta oscillation detection in similar fashion to prior studies^23,84^. We applied a zero-phase shift 4-8 Hz bandpass filter to the single-channel (cortical contact) LFP data. We set a channel-specific cutoff at 3 standard deviations above the mean and was applied to each channel’s Hilbert envelope to identify peaks of theta oscillations. We identified the start and stop of each theta oscillation event as when the power dropped below 1 standard deviation above the channel mean. Theta bursts were included for analysis if their durations fell within 375 and 1000ms (defined as 3 cycles of 8 Hz and 3 Hz, respectively).

### Peri-event time histograms

To construct peri-event time histograms across experimental conditions, we used a bin width advised by Scott’s optimization method, which optimizes bin size to event density^85^. For PETH locked to sentence onset, we assessed all ripples in the −0.5 to 10 second window relative to onset of word list presentation and utilized an optimized bin size of 200ms with 7-point smoothing. To determine significant time bins, 5000 iterations of peri-presentation onset time histograms were computed by circularly jittering ripple times across the two-second pre-word list presentation window. This enabled us to maintain statistical power despite a low number of ripple events. We performed a permutation test determine significant time windows. Time windows that were in the top or bottom 2.5% of the distribution (thus implementing an alpha-value of 0.05) were denoted as significant. A similar approach was taken for the peri-verbal recall PETH, but here, statistical power was computed by circularly jittering ripple times 5000 times across the same −4 to 1 second window relative to onset of verbal free recall (while implementing 90ms bin size with 7-point smoothing). Finally, to compare PETH around the onset of effective vs poor recall periods, we assessed the same −4 to 1 second window with identical 230ms and 4-point smoothing for the two PETH. To determine significance, we shuffled labels of effective and poor recall and generated a distribution of mean SWR rate differences (5000 iterations). This generated a distribution by which we identified significant timebins.

### Cross-correlograms

To examine “co-rippling” between two brain regions, we used the ripple peak time index to produce cross-correlograms. For each cortical electrode, we computed a cross-correlation between detected cortical ripples and SWRs. We pooled these cross-correlations across trials for each electrode pair in each participant, which creates a single cross-correlogram for each pair of contacts. We normalized the amplitude of each cross-correlation by the duration of the time window of interest and number of electrode pairs such that values represented coincidences per second. For this analysis, we implemented 10ms bins. To determine significance of synchronous ripple events, we jittered cortical ripple event timing and constructed a new cross-correlogram. We repeated this procedure 5000 times and used to generate a null distribution for a permutation test. We applied this procedure separately for ATL contacts and semantic network contacts.

### Temporal resolution of coincident ripple events

To determine the temporal resolution of co-ripple events relative to verbal recall onset, we implemented proportions to normalize for the increase in SWR events that occurs prior to verbal free recall. That is, instead of quantifying the rate of coincident ripple events, we quantified the chance that a SWR event was coincident with a coincident ATL cortical ripple event prior to verbal free recall. We defined a coincident ripple event as two ripple events where the peak-to-peak duration was less than 100ms (defined as half of the 200ms maximum ripple duration), as the peak of ripple events are more readily detectable compared to the onset/offset (which differs depending on manually-defined thresholds). We constructed a PETH for the −4 to 2 second window relative to verbal recall onset implementing 300ms bins with 8-point smoothing. To determine significance across timebins, we jittered cortical ripple event timing and constructed a new PETH. We repeated this procedure 5000 times and to create a null distribution for a permutation test.

## Acknowledgements

We would like to thank the subjects for volunteering their time and effort to participate in our study. We also thank the Epilepsy Monitoring Unit staff at the North Shore University Hospital for their support throughout the study. We thank Yitzhak Norman and Jose Herrero for providing helpful feedback on the analysis and methods. This work was supported by NIH grant R01DC019979 (to S.B.).

## Authorship statement

AMishra, SA, and SB designed the study. AMehta performed clinical electrode implantation. AMishra, SA, EE, NM, and EF conducted the experiments. AMishra and SA conducted all data analysis. AMishra and SB prepared the initial draft of the manuscript. All authors approved of the final manuscript draft.

