## Supplementary Material for "Hippocampal sharp wave ripples and coincident cortical ripples orchestrate human semantic networks"

**Table 1.** Demographic information for n=9 participants, including number of semantic network and anterior temporal lobe contacts implemented in analysis.

| Subject Number | Age (years) | Primary Language | # of ATL electrodes | # of Sentence-Responsive Contacts | # of Non-Sentence-Responsive Contacts | Mean # of Words Recalled per Trial |
|---|---|---|---|---|---|---|
| 1 | 28 | English | 3 | 28 | 2 | 3.3 |
| 2 | 35 | English | 13 | 11 | 10 | 6.2 |
| 3 | 19 | English | 36 | 5 | 5 | 5.5 |
| 4 | 29 | English | 70 | 28 | 19 | 3.0 |
| 5 | 36 | English | 13 | 8 | 0 | 5.4 |
| 6 | 58 | English | 2 | 2 | 15 | 5.1 |
| 7 | 19 | Spanish | 32 | 12 | 11 | 3.0 |
| 8 | 32 | English | 14 | 37 | 9 | 3.9 |
| 9 | 55 | English | 16 | 19 | 8 | 5.1 |

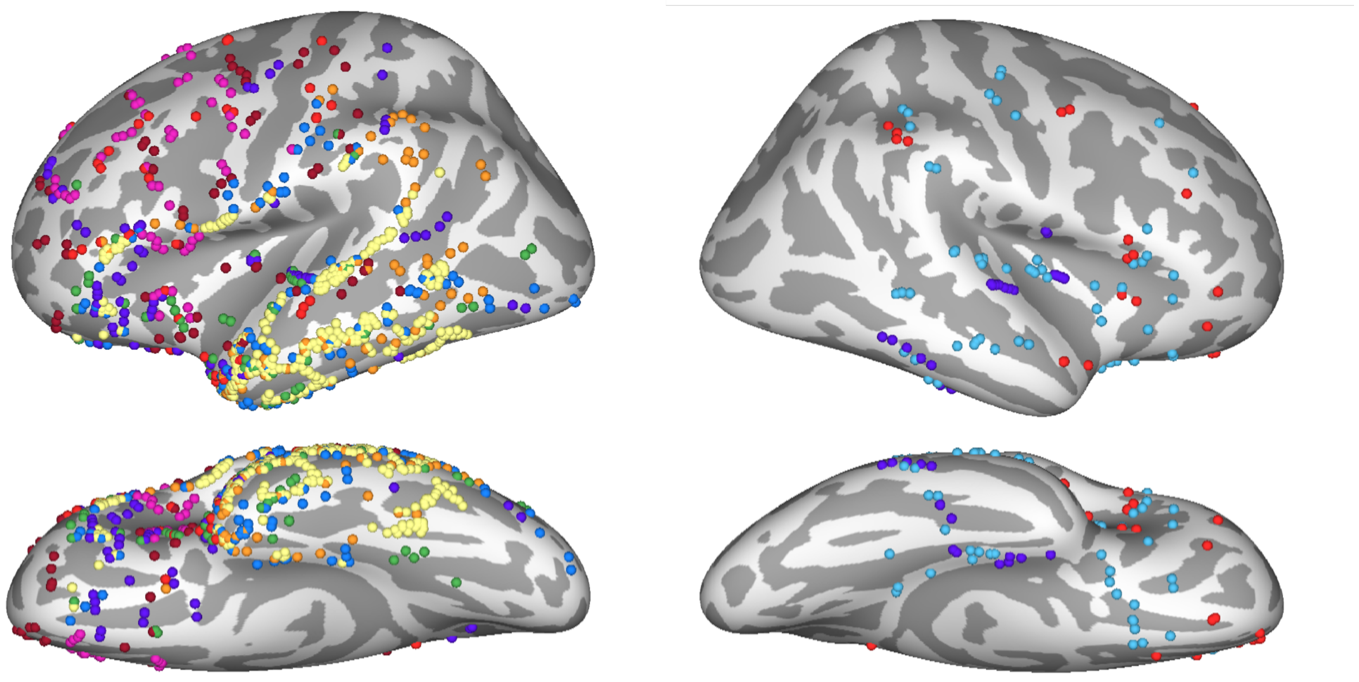

**Figure S1.** Electrode coverage across subjects depicted on an inflated brain (in lateral and inferior views of left and right hemispheres). Each color represents the set of cortical contacts from one unique patient.

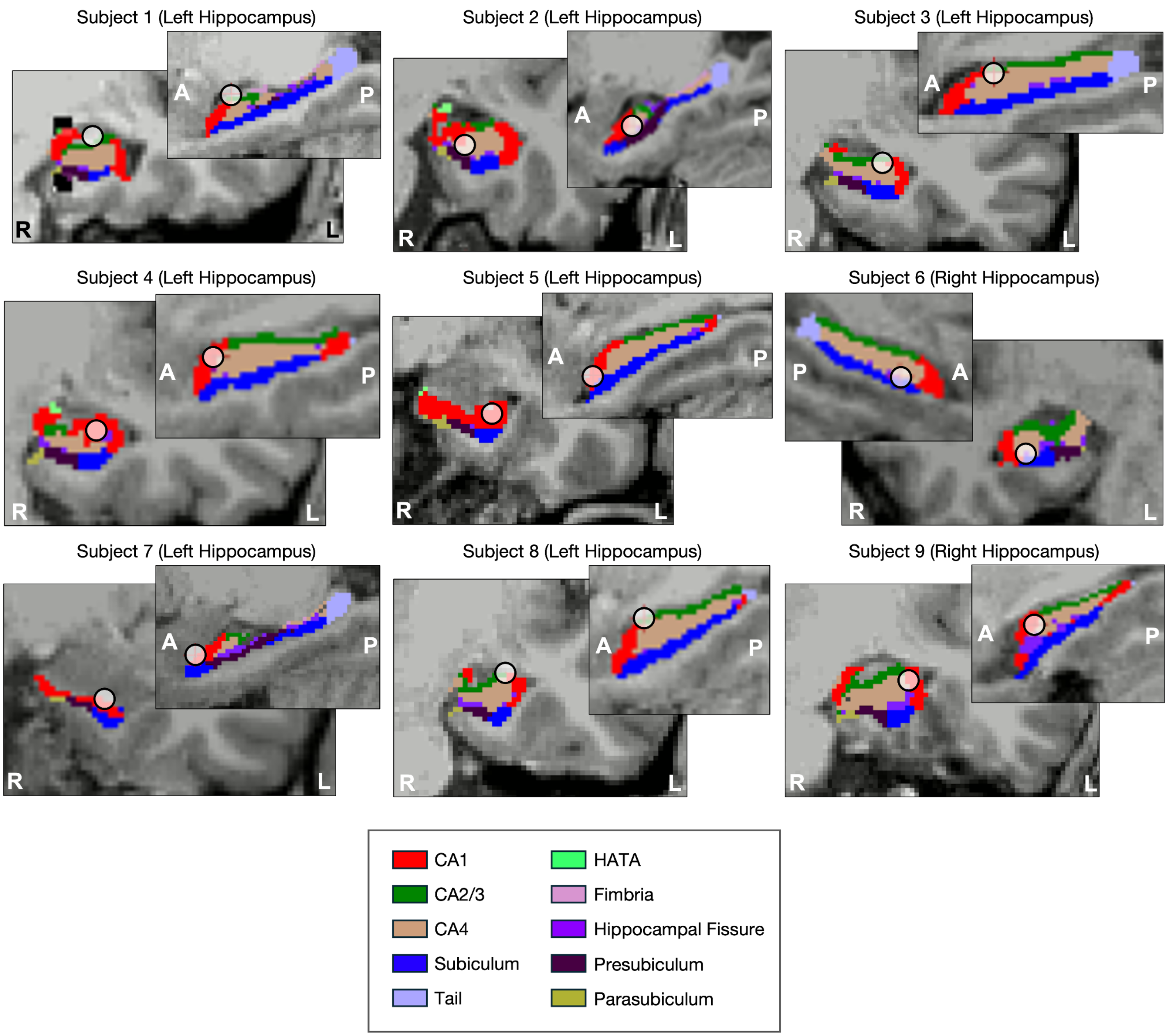


**Figure S2.** Hippocampal parcellations for each patient, in subject space, with white translucent marker indicating selected hippocampal contact and reference contact in coronal (background) and sagittal (inset) views.

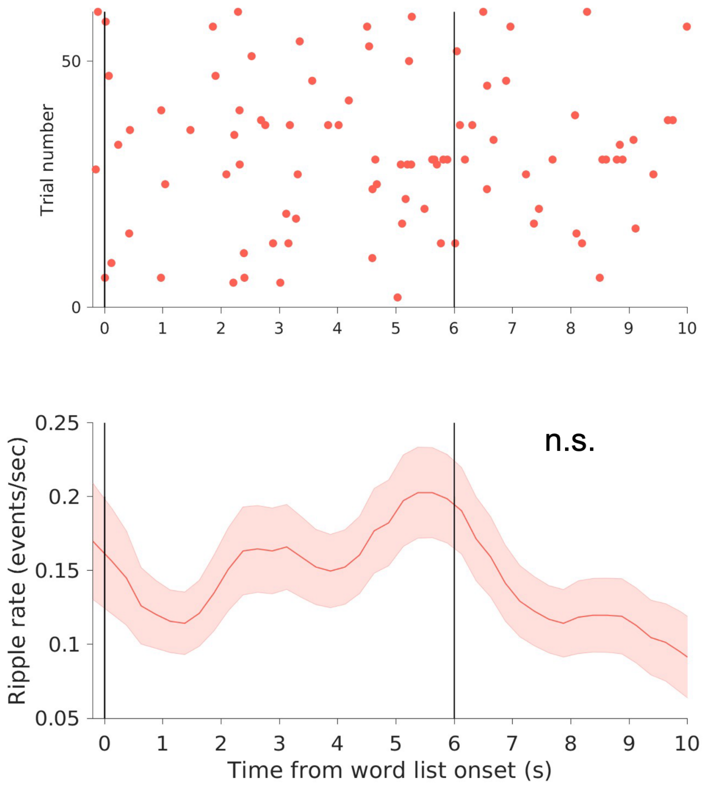


**Figure S3.** SWR raster plot and peri-event time histogram time-locked to the onset of word list presentation for two subjects (n=60 trials) with discrepancy in word list presentation length. In these patients, word presentation was 0.5 Hz (rather than 0.67 Hz) such that presentation of a set of 12 words was a total of 6 seconds in duration. Shaded areas represent one bootstrap standard error computed over SWR events. n.s., not significant (permutation test compared to shuffled SWR times in the 2-second post-trial resting period)

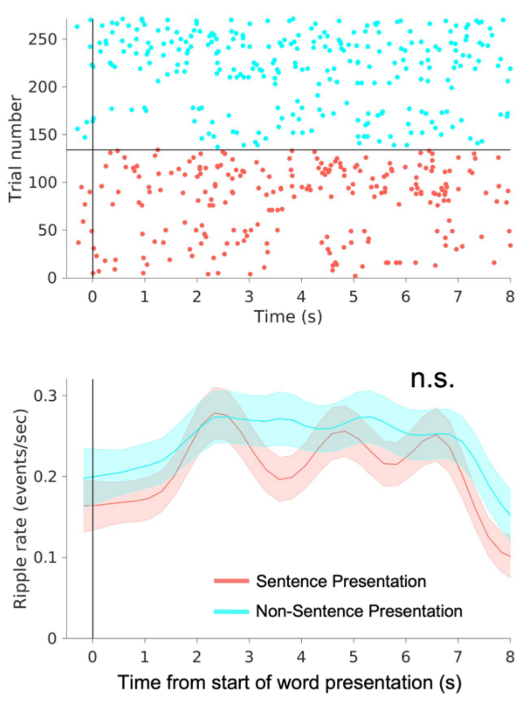


**Figure S4.** SWR raster plot and peri-event time histogram time-locked to the onset of word list presentation comparing rate of SWR for sentence (n=134 trials, red) and non-sentence (n=137 trials, blue) word lists. Shaded areas represent one bootstrap standard error computed over SWR events for each trial condition. n.s., not significant (permutation test shuffling trial labels)

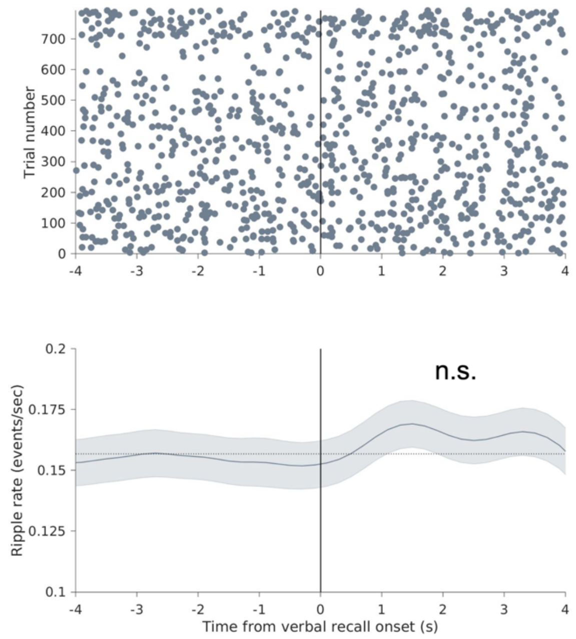


**Figure S5.** SWR raster plot and peri-event time histogram time-locked to the onset of patient vocalizations during the task block that exceeded 6SD relative to baseline indicating no significant relationship between random vocalizations and SWRs. Shaded areas represent one bootstrap standard error computed over SWR events. n.s., not significant (permutation test shuffling SWR times across this PETH epoch)

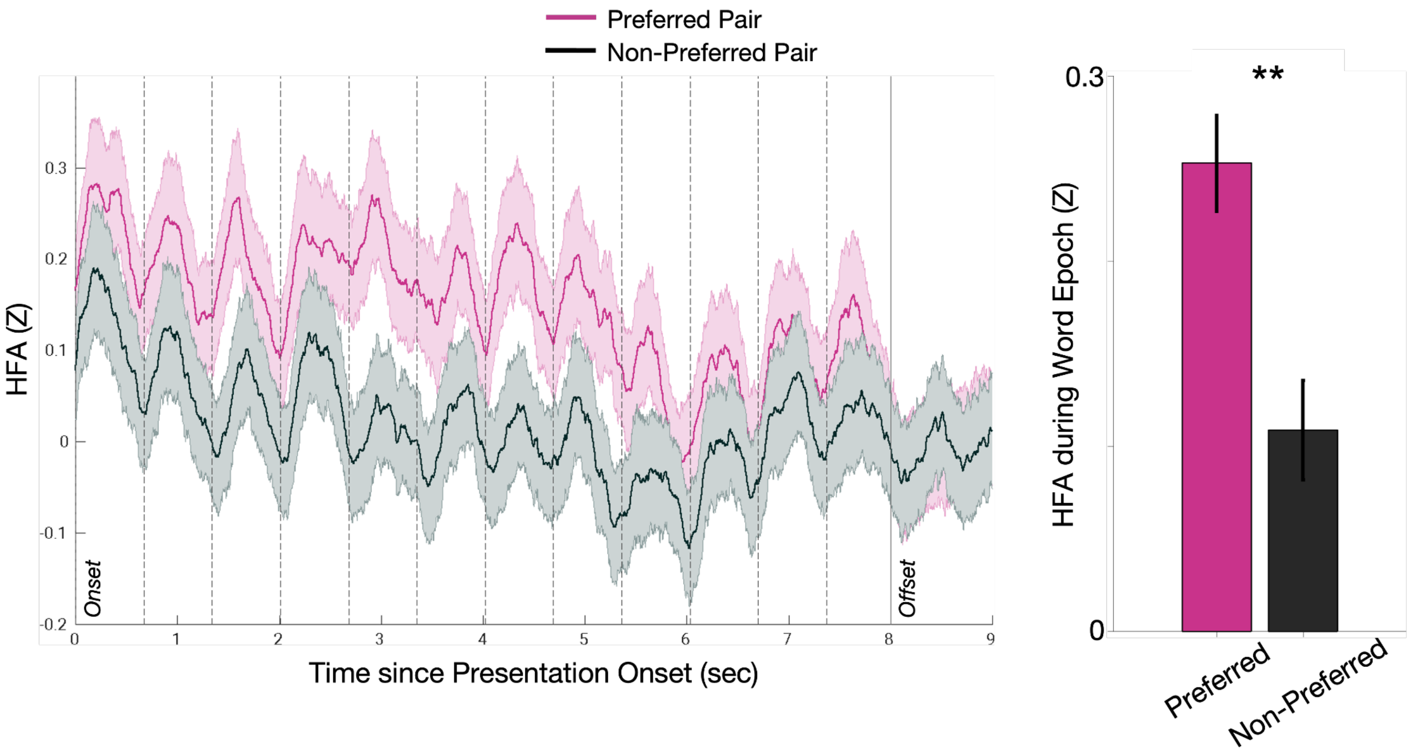


**Figure S6**. (Left) Magnitude of HFA during presentation of preferred (congruence between trial type and contact preference) vs non-preferred contact-trial pairings (non-congruence between trial type and contact preference) for n=212 contacts across 7 subjects (n=30 trials per subject, n=15 sentence and n=15 word list) with a word list presentation duration of 8 seconds. (Two subjects were removed from this analysis due to a difference in duration of presentation.) Presentation of words was followed by a 2-second baseline period. Dashed vertical lines represent the point of word onset. Note that the congruent trial-contact pairs exhibited increased HFA relative to non-congruent pairs and the difference between the two diminishes after offset of word list presentation. (Right) Magnitude of HFA activation during presentation of words (500ms epochs) for preferred compared to non-preferred trials for n=9 subjects. Error bars represent one standard error across contacts. The median Z-scored HFA for preferred and non-preferred word list perception were significantly different (0.226 compared to 0.091; Z=5.056, p<0.001). Stars represent significance at p<0.05.

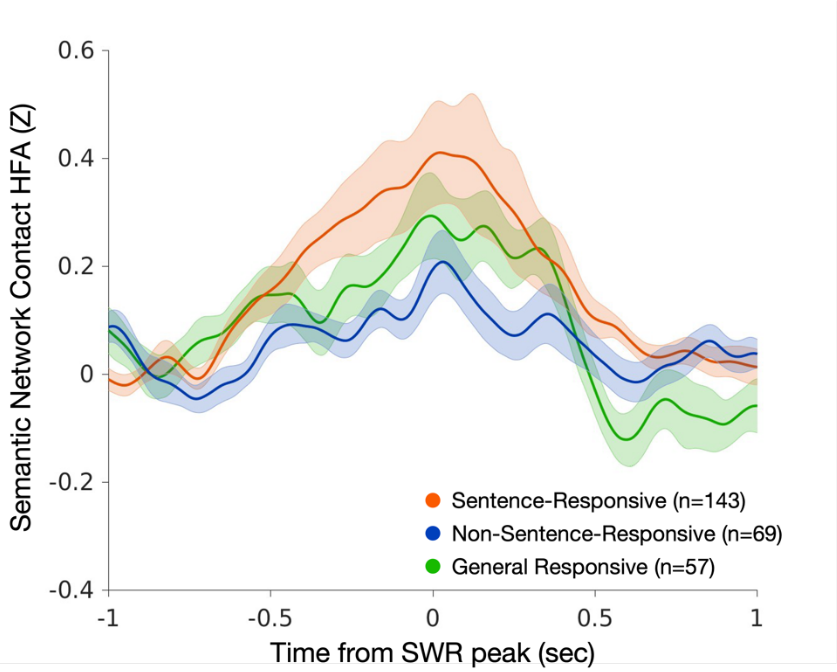


**Figure S7.** SWR peak-locked HFA in sentence-preferential (n=143 contacts), non-sentence-preferential (n=69 contacts), and general language-responsive contact (n=57 contacts) for SWR events associated with verbal free recall. Shaded area represents one standard error across contacts.

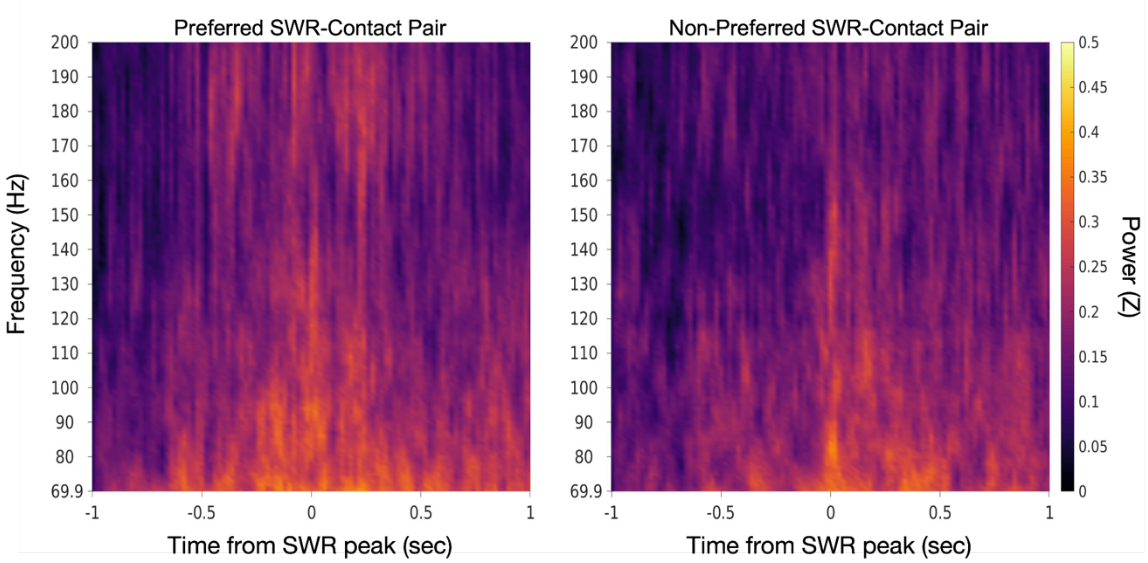


**Figure S8.** Spectrograms of response SWR-locked high frequency activity in preferred (left) versus non-preferred (right) trial-contact alignments. Increase in HFA is found to be broadband.

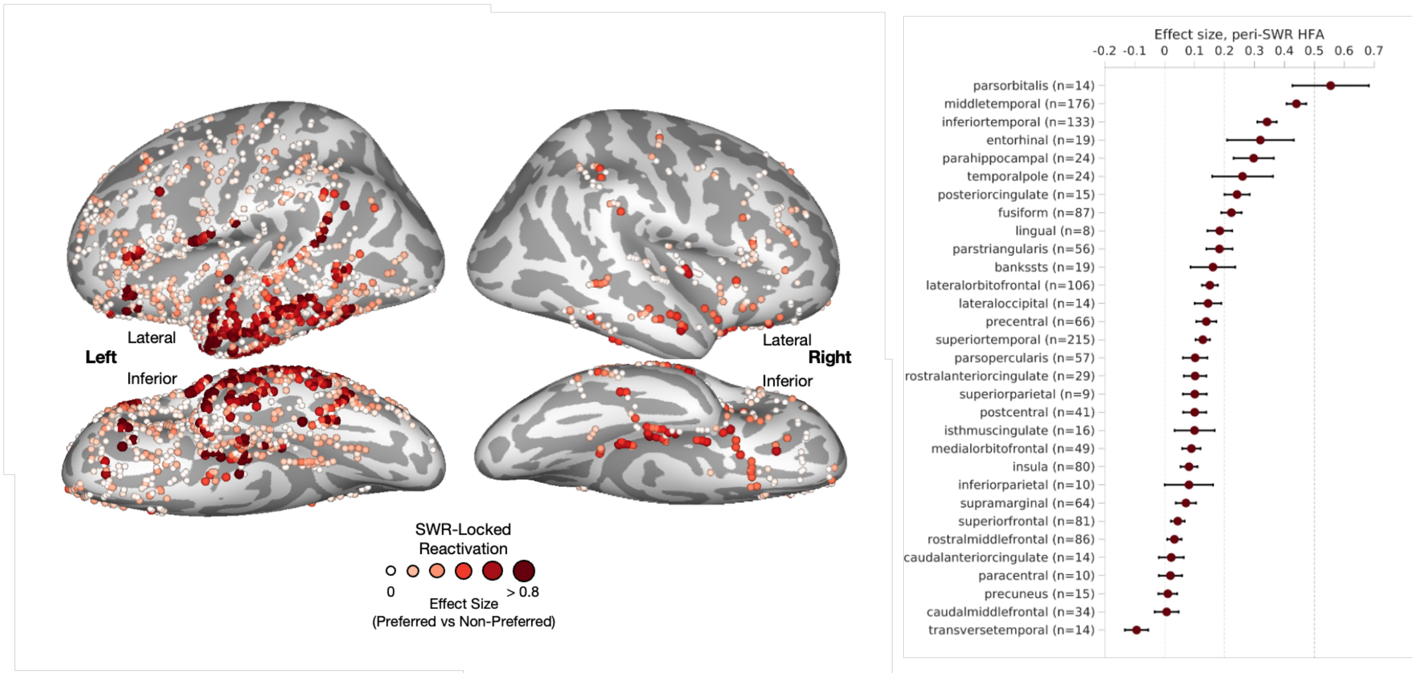


**Figure S9**. (Left) Effect size (in Cohen's d) of HFA increase associated with SWR during the recall period compared to a matched pre-SWR baseline period across all cortical contacts across subjects. Darker colors represent increased effect size (SWR-locked HFA in 500ms window around SWR peak compared to matched duration in pre-SWR baseline). (Right) Mean effect size by DK atlas region (with number of contacts present in the parcel), error bars represent one SEM. Dashed lines are added to distinguish medium (0.5-0.8) and small (0.2-0.5) effect sizes.

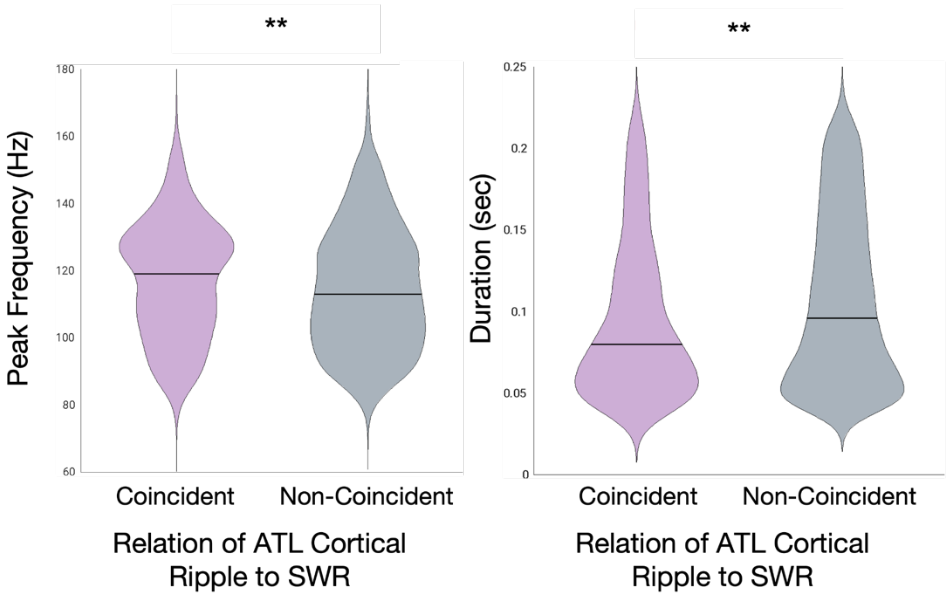


**Figure S10**. Distribution of (A) peak cortical ripple frequency and (B) cortical ripple duration for n=1187 coincident cortical ripple events and n=14762 non-coincident cortical ripple events occurring within the recall period. Black line represents group median. Stars denote $p<0.05$ (Mann-Whitney U test). Coincident ripples are found to be shorter and of higher frequency compared to non-coincident ripples.

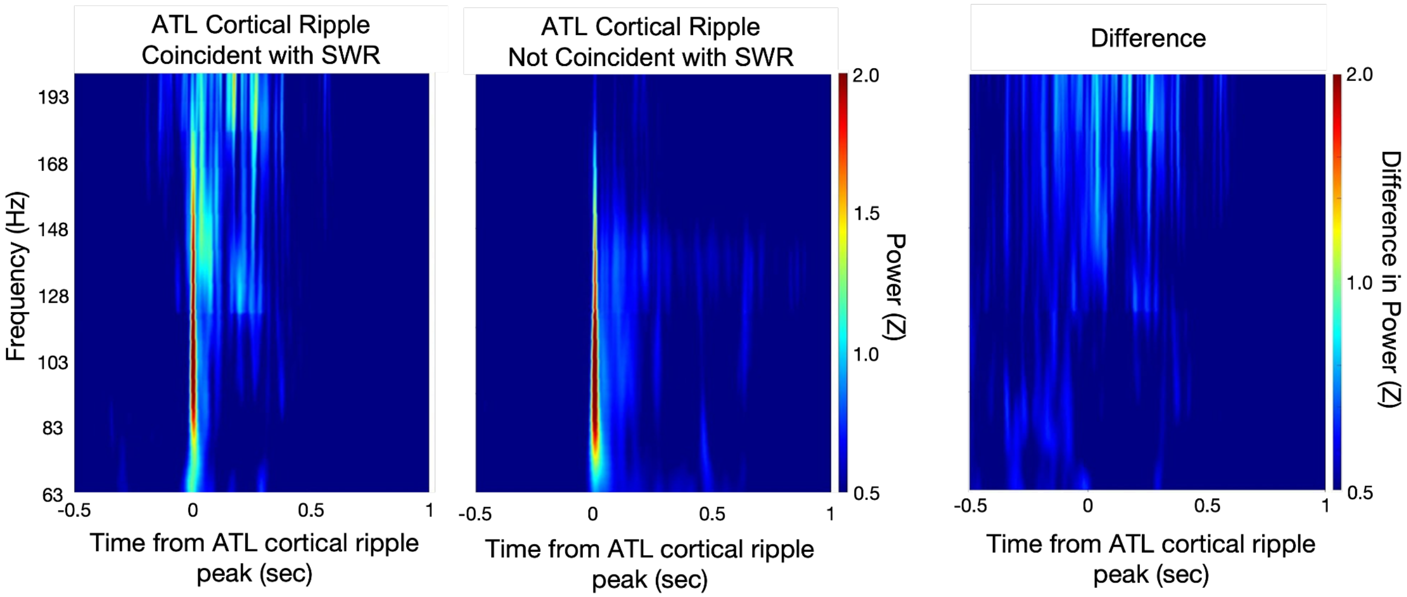


**Figure S11**. Comparison of high-frequency spectral profiles locked to peak of (A) n=1187 coincident cortical ripple events and (B) n=14762 non-coincident cortical ripple events occurring within the recall period. (C) Represents the difference in power spectra. Warmer colors indicate a higher spectral power in that frequency-time combination.

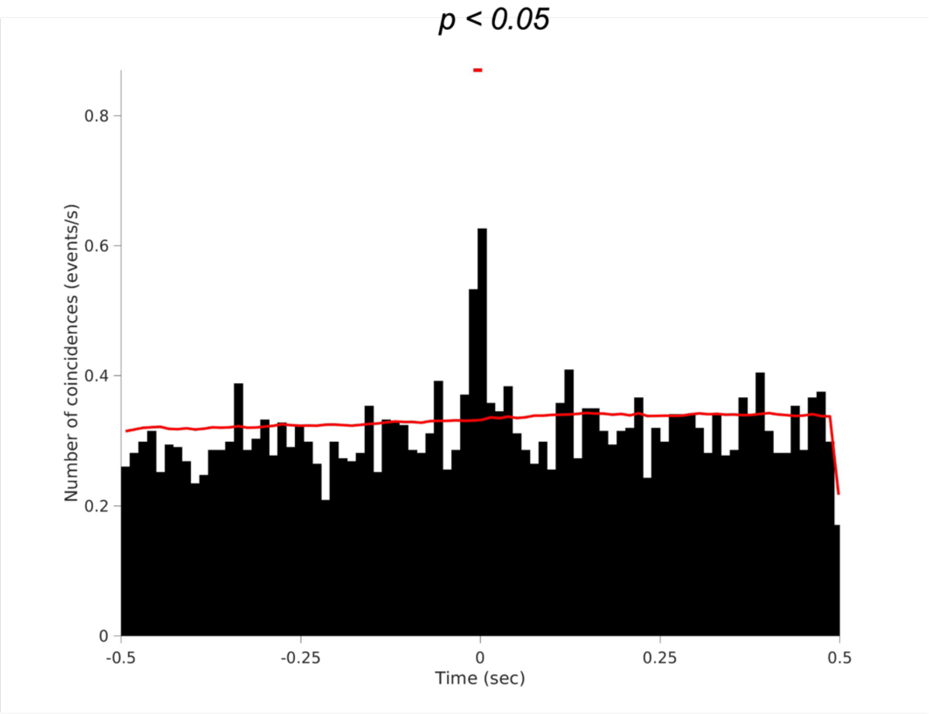


**Figure S12.** Cross-correlograms between semantic network cortical ripple events and SWR events within the recall period across n=9 subjects. Red line denotes significant time bins at p<0.05 (permutation test with jittered cortical ripple timing), indicating high coincidence.

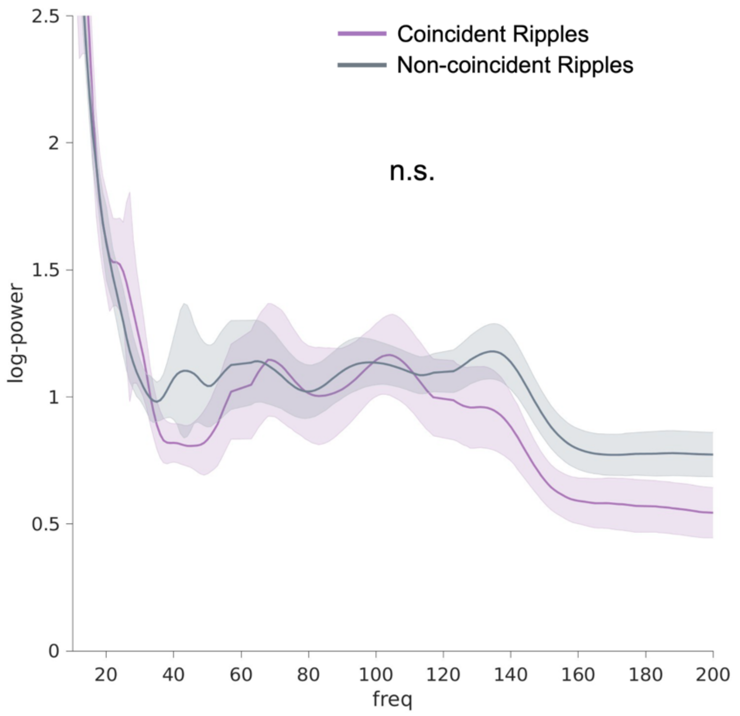


**Figure S13.** Spectral profile (1/f plot) of semantic network cortical ripples associated with verbal recall by whether the cortical ripple was associated with a SWR event compared to not. n.s., not significant. (cluster-based permutation test clustering frequencies)
